# Integrative tissue-specific functional annotations in the human genome provide novel insights on many complex traits and improve signal prioritization in genome wide association studies

**DOI:** 10.1101/028464

**Authors:** Qiongshi Lu, Ryan L Powles, Qian Wang, Beixin J He, Hongyu Zhao

## Abstract

Extensive efforts have been made to understand genomic function through both experimental and computational approaches, yet proper annotation still remains challenging, especially in non-coding regions. In this manuscript, we introduce GenoSkyline, an unsupervised learning framework to predict tissue-specific functional regions through integrating high-throughput epigenetic annotations. GenoSkyline successfully identified a variety of non-coding regulatory machinery including enhancers, regulatory miRNA, and hypomethylated transposable elements in extensive case studies. Integrative analysis of GenoSkyline annotations and results from genome-wide association studies (GWAS) led to novel biological insights on the etiologies of a number of human complex traits. We also explored using tissue-specific functional annotations to prioritize GWAS signals and predict relevant tissue types for each risk locus. Brain and blood-specific annotations led to better prioritization performance for schizophrenia than standard GWAS p-values and non-tissue-specific annotations. As for coronary artery disease, heart-specific functional regions was highly enriched of GWAS signals, but previously identified risk loci were found to be most functional in other tissues, suggesting a substantial proportion of still undetected heart-related loci. In summary, GenoSkyline annotations can guide genetic studies at multiple resolutions and provide valuable insights in understanding complex diseases. GenoSkyline is available at http://genocanyon.med.yale.edu/GenoSkyline.

## Introduction

Functionally annotating the human genome is a major goal in human genetics research. After years of community efforts, a variety of experimental and computational approaches have been developed and applied for genomic functional annotation. Comparative genomics studies have shown that approximately 4.5% of the human genome is conserved across mammals^1^. Furthermore, the rich collection of epigenomic data generated by large consortia (e.g. ENCODE^2^ and Epigenomics Roadmap Project^3^) also provides great insight for understanding the functional effects of the genome, especially in terms of non-coding regulatory machinery. To best utilize these rich data, we recently developed GenoCanyon^4^, a non-coding functional prediction approach based on integrative analysis of annotation data, whose performance was demonstrated through predicting well-studied regulatory DNA elements. GenoCanyon provides general predictions of non-coding functional regions in the human genome but does not fully utilize cell-type-specific information of epigenomic data. Incorporating cell-type-specific or tissue-specific information into annotation tools is essential not only for understanding the basic biology of the genome, but also for better characterizing genetic variation, as in the functional interpretation of risk loci identified from genome-wide association studies (GWAS).

GWAS has been a great success in the past decade, yet challenges still remain in both identifying additional risk variants and interpreting GWAS results. Current practice employs a significance threshold (i.e. 5×10^−8^) that controls family-wise error rate. Yet this approach is known to be underpowered when effect sizes are weak or moderate at risk loci^5^. Moreover, nearly 90% of the genome-wide significant hits in published GWAS are located in non-coding regions whose functional impact to human complex traits is largely unknown^6^. Complex linkage disequilibrium (LD) patterns also hinder our ability to identify real functional sites among correlated SNPs. Several methods have been proposed to integrate annotation data for better prioritizing GWAS signals and their effectiveness has also been well demonstrated^7-10^. Tissue-specific functional annotations have the potential to bring even more biological insights to post-GWAS analysis and help understand complex disease etiology.

In this paper, we introduce GenoSkyline, a tissue-specific functional prediction tool based on integrated analysis of epigenomic annotation data. We demonstrate its ability to identify tissue-specific functionality from its performance to rediscover a number of experimentally validated non-coding elements. Next, we show valuable biological insights GenoSkyline can provide in post-GWAS analysis through integrative analysis of 15 human complex traits. We believe that GenoSkyline will prove to be a powerful tool for human genetics research because of its abilities to assess tissue-specific enrichment of GWAS signals, better prioritize GWAS signals, and offer biological interpretations of risk loci.

## Results

### Predicting tissue-specific functional regions in the human genome

The posterior probability of being functional given the annotation data is used to measure tissue-specific functional potential of each nucleotide in the human genome (**Online Methods**). It will be referred to as GenoSkyline (GS) score in following sections. We calculated GS scores for 7 human tissue types; brain, gastrointestinal tract (GI), lung, heart, blood, muscle, and epithelium (**Supplementary Table 1**). With a GS score cutoff of 0.5, 22.2% of the human genome is predicted to be functional in at least one of these tissue types, while 1.7% is functional in all 7 tissues (**Figure 1a**). Since GS score has a bimodal pattern, these results are not sensitive to cutoff choice (**Supplementary Notes**).

**Figure 1.**
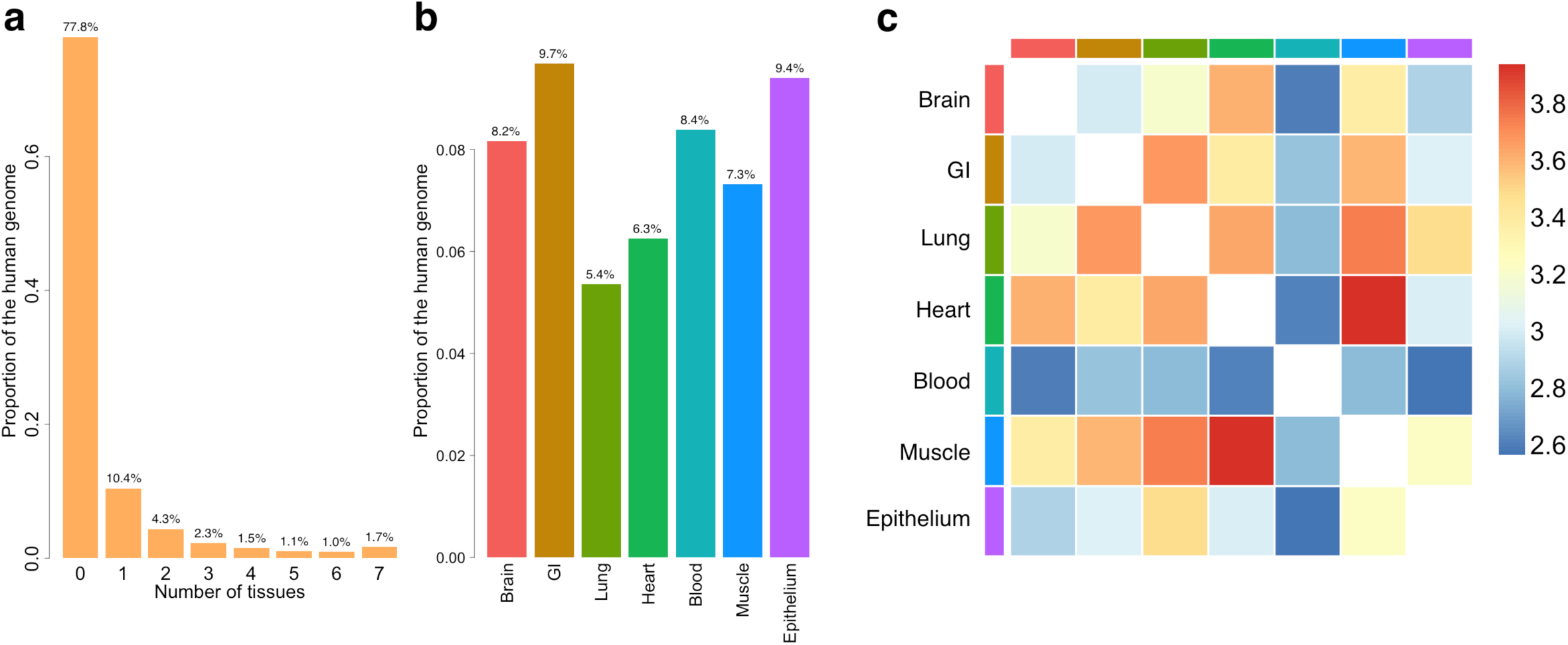
General characteristics of GenoSkyline annotations. (a) Number of tissues in which nucleotides are functional. (b) Proportion of functional genome for each tissue type. (c) Overlap of functional regions across seven tissue types. The scale is log odds ratio.

Across tissue types, the percentage of predicted functional genome ranges from 5.4% (Lung) to 9.7% (GI) (**Figure 1b** and **Supplementary Table 2**). The overlap between heart-specific and muscle-specific functional regions is the largest among all pairs of tissues. Interestingly, although the percentage of functional genome in blood (8.4%) is similar to other tissue types, it overlaps less with the functional regions in other tissues (**Figure 1c**). This is consistent with the recent discovery that blood has the lowest levels of eQTL sharing with other tissues^11^.

### Investigating the performance of tissue-specific functional annotations Beta-globin gene complex

We now demonstrate GenoSkyline’s ability to predict tissue-specific functionality using a variety of experimentally validated functional machinery. Beta-globin (*HBB*) gene complex is an extensively studied genomic region on chromosome 11, containing 6 genes and 23 cis-regulatory modules (CRMs) that are known to control both the timing and the spatial pattern of gene expression^12,13^. We compared GS scores for different tissue types in this region. Not surprisingly, blood-specific functionality was observed (**Figure 2a**). Among the 6 genes in this region, adult globin genes *HBB* and *HBD,* as well as pseudogene *HBBP1* are captured well by blood-specific GS scores (**Supplementary Table 3**). However, embryonically expressed *HBE1,* fetally expressed *HBG1* and *HBG2,* and the CRMs that regulate these genes have lower GS scores. This is possibly because 18 of the 24 cell lines used for developing blood-specific GS scores were acquired from adult samples (**Supplementary Table 1**). The mean blood-specific GS score in these genes increases from 0.388 to 0.704 after removing *HBG1, HBG2,* and *HBE1.* Similarly, a substantial boost in mean GS score is observed after removing CRMs regulating the embryonic and fetal globin genes (**Figure 2b**, **Supplementary Figure 1**). Compared with GenoCanyon, GenoSkyline provides less sensitive but highly specific functional predictions. Its ability of identifying tissue-specific functional coding and non-coding DNA elements has the potential to benefit diverse types of biological studies.

**Figure 2.**
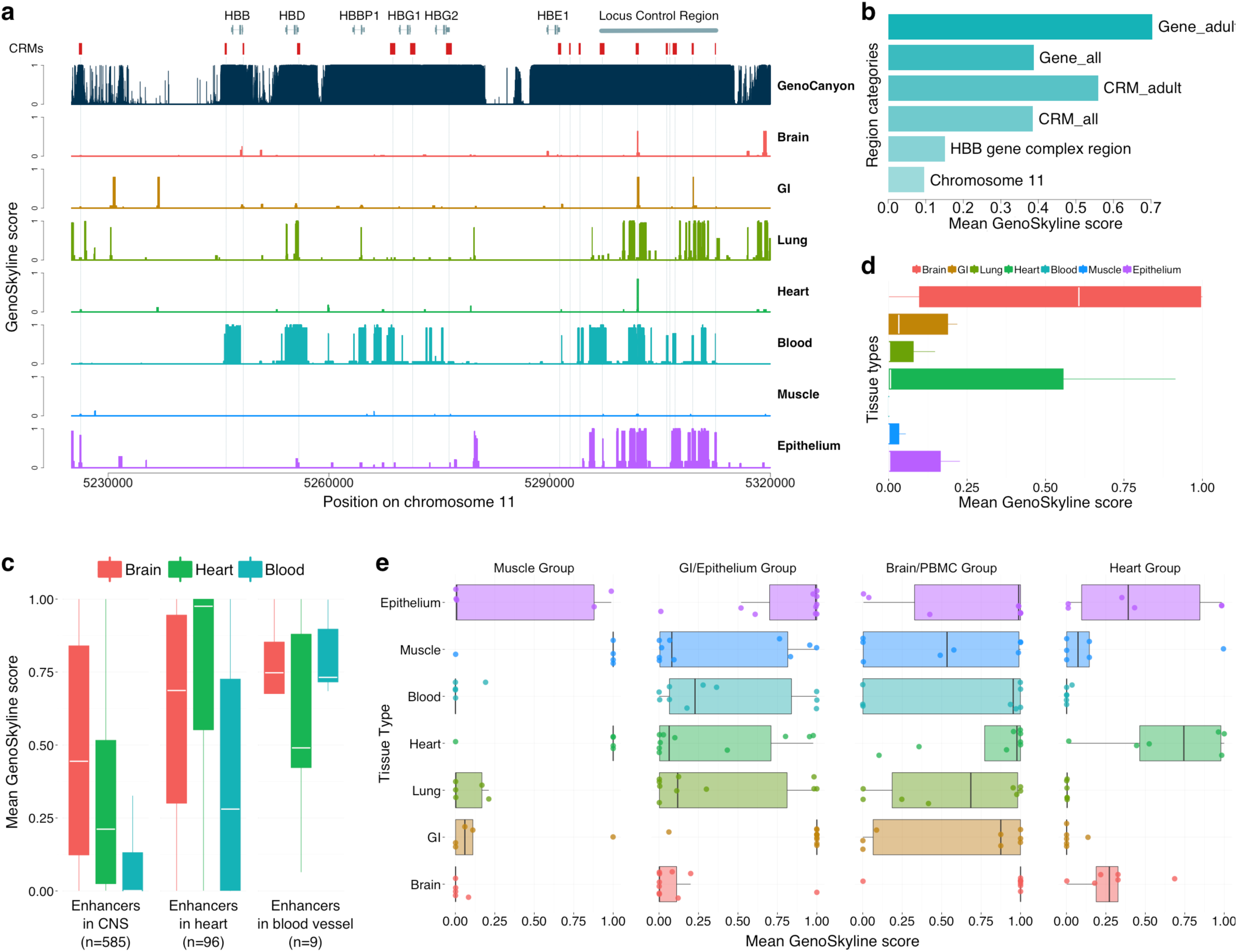
Case studies of *HBB* gene complex, *in vivo* enhancers, and regulatory miRNAs. (a) Comparison of GenoCanyon prediction and GenoSkyline scores for seven tissues in *HBB* gene complex region. Red boxes mark the locations of CRMs. The number of red boxes is less than 23 because some CRMs are next to each other. (b) Mean blood-specific GS score for different region categories. (c) Boxplot of mean GS scores for enhancers in CNS, heart, and blood vessel. (d) Boxplot of mean GS scores for 11 human-accelerated elements near *NPAS3.* (e) Boxplot of mean GS scores for tissue-specific regulatory miRNAs.

### Tissue-specific enhancers

*In vivo* enhancers with tissue-specific activity in central nervous system (CNS; n=585), heart (n=96), and blood vessel (n=9) were downloaded from VISTA enhancer browser^14^ (**Online Methods**). Mean GS scores for brain, heart, and blood tissues were calculated for each enhancer. Brain-specific and heart-specific GS scores were substantially higher in their respective enhancer categories compared to GS scores of non-relevant tissue types. Additionally, the mean blood-specific GS score also stands out for enhancers with observed activity in blood vessel despite the limited sample size (**Figure 2c**). In a separate study, 11 human-accelerated elements near the brain developmental transcription factor *NPAS3* have been identified to act as tissue-specific enhancers within the nervous system^15^. Brain-specific GS scores for these enhancers are substantially higher than those for other tissue types (**Figure 2d**), concurrent with previous results.

### Regulatory miRNAs

Next, we test if GenoSkyline could also capture miRNAs expressed exclusively in certain tissue types. Liang et al. studied the tissue specific expression pattern of eight groups of miRNAs^16^. We extracted and annotated four groups (groups I, II, III+IVa, and V from Liang et al.) that could be represented by the currently available tissue types in GenoSkyline annotations. These four groups of miRNAs were found to be expressed preferentially in skeletal/cardiac muscle, organs lined with epithelium, brain/peripheral blood mononuclear cell (PBMC), and heart, respectively through unsupervised clustering. The most relevant tissue types suggested by GenoSkyline for these four groups are muscle/heart, GI/epithelium, brain, and heart, respectively (**Figure 2e**). Our results based on integrative analysis of epigenomic data are consistent with the tissue-specific expression pattern reported by Liang et al.

### Inter-genic regulation of myosin heavy chain

We applied GenoSkyline to a validated biologic switch in cardiac development and disease. Myosin heavy chain (MHC) is the major contractile protein in human striated muscle^17^. Cardiac muscle cells primarily express two isoforms, alpha-MHC (*MYH6*) and beta-MHC (*MYH7*)^18^. The ratio of alpha-to-beta isoforms determines cardiac contractility and allows for effective response to a wide range of physiologic and pathologic stimuli^19^. Alpha-to-beta ratio decreases in cardiac diseased states^20,21^, and reversal of this shift is associated with better clinical outcomes^22^. miRNAs can regulate alpha-to-beta isoform shift, and prior studies in rodents have outlined a network of crosstalk between intronically expressed miRNAs and their host muscle genes^23,24^. For instance, mir-208a, on an intron of *MYH6*, is a positive regulator of beta-MHC by targeting transcription factors that repress its expression^24^. GS scores for *MYH6* and mir-208a accurately reflect their cardiac-specific expression, whereas *MYH7* and mir-208b exhibit strong signals in both skeletal and cardiac tissue (**Figure 3a** and **Supplementary Table 4**). This corresponds to known expression pattern of *MYH7* and mir-208b in slow twitch skeletal muscle fibers^17^ as well as heart. We also explored tissue-specific functionality of two known distal enhancers of mir-208b identified on VISTA Enhancer Browser, hs2330 and hs1670. GS scores for hs2330 mirror *MYH7*/mir-208b signals. Interestingly, GS scores for hs1670, a distal enhancer flanking mir-208b, are also strong in nervous and GI tissue, a finding that agrees with its observed expression pattern in other tissues (based on VISTA Enhancer Browser data). Collectively, these results show that GenoSkyline can replicate the tissue-specific expression pattern of a complex inter-gene regulatory network.

**Figure 3.**
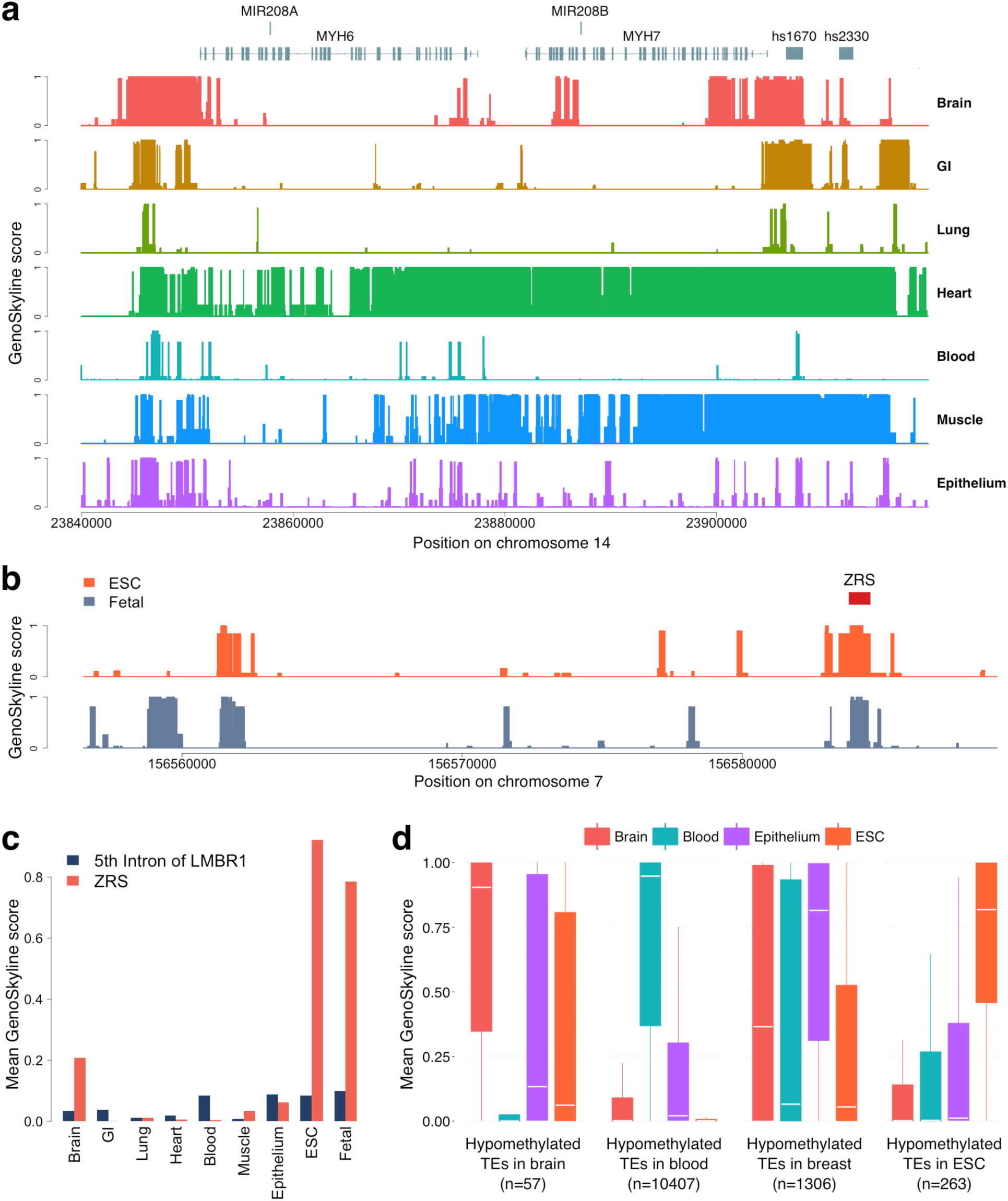
Case studies of MHC, ZRS, and hypomethylated TEs. (a) GenoSkyline scores for seven tissues in the genomic region surrounding *MYH6* and *MYH7.* (b) ESC-specific and fetal-cell-specific GS scores for the 5th intron of *LMBR1*. The red box marks the location of ZRS. (c) Bar plot of the mean GS scores for the 5th intron of *LMBR1* and ZRS across nine tissue and cell types. (d) Boxplot of mean GS scores for four groups of hypomethylated TEs.

### Zone of polarizing activity regulatory sequence

GenoSkyline can also be generalized to identify tissue specificity outside of the 7 core categories discussed here, based on available experimental data. For example, Zone of polarizing activity regulatory sequence (ZRS), a well-studied developmental enhancer, is located in the fifth intron of *LMBR1* gene. Acting as an enhancer of *SHH,* ZRS has been shown to play a crucial role in limb development^25^. However, none of the seven tissue types in GenoSkyline suggest ZRS’s functionality (**Supplementary Figure 2**). In order to see if ZRS could be identified using epigenomic data of other cell types, we extended GenoSkyline to two new groups of cells that are potentially important for development, embryonic stem cells (ESC) and fetal cells (**Supplementary Table 5**). Both ESC and fetal-cell-specific GS scores successfully identified ZRS with high resolution (**Figures 3b and 3c**). This example shows that GenoSkyline is a flexible framework. Researchers could develop their own cell-group-specific functional annotations if ChIP-seq data are available for the cells of interest.

### Hypomethylated transposable elements

A recent study of genome-wide DNA methylation status identified tissue-specific hypomethylated transposable elements (TE) exhibiting enhancer activities^26^. We downloaded four groups of TEs that are hypomethylated in ESC H1, fetal brain/primary neural progenitor cells, adult breast epithelial cells, and PBMC/adult immune cells, respectively (**Online Methods**). Although DNA methylation data were not used for developing GenoSkyline, we were still able to provide highly consistent results, suggesting tissue-specific functionality of these TEs in ESC, brain, epithelium, and blood cell, respectively (**Figure 3d**).

### Analyzing tissue-specific enrichment for 15 human complex traits

In the sections above, we demonstrated GenoSkyline’s ability of identifying tissue-specific functional regions in the human genome. Next, we focus on how GenoSkyline could help us understand human complex traits. Finucane et al. recently proposed using LD score regression to partition heritability of complex traits by functional categories^27^. We applied LD score regression on 15 human complex diseases and traits (**Supplementary Table 6**), and calculated the tissue-specific enrichments using GenoSkyline annotations (**Online Methods**).

Our analysis successfully replicated some well-known findings and also provided novel insights to these complex traits (**Figure 4**; **Supplementary Figure 3**). For schizophrenia, enrichment in brain is much stronger than in any other tissue type (*p* = 6.52×10^−26^), while highly significant enrichment could be observed in heart (*p* = 2.30×10^−7^) and blood (*p* = 1.65×10^−5^) as well. Brain is also the most enriched tissue for anorexia nervosa (*p* = 4.86×10^−2^) despite the substantially weaker signal. For three autoimmune diseases (Crohn’s disease, ulcerative colitis, and rheumatoid arthritis), the strongest enrichment was in blood. However, solid enrichment in GI could also be observed for both Crohn’s disease and ulcerative colitis, but not rheumatoid arthritis. Sex-stratified summary statistics were available for two anthropometric traits - body mass index (BMI) and waist-hip ratio (WHR) adjusted for BMI^28,29^. Therefore, we performed gender-specific analyses for these two traits. Consistent with recently published results^27^, brain possesses the strongest enrichment for BMI. Interestingly, the enrichment in brain is stronger in female samples (*p* = 1.21×10^−8^) than in male samples (*p* = 2.39×10^−6^), while epithelial tissue may play a more important functional role in male samples (*p* = 1.34×10^−3^ in males and 2.83×10^−2^ in females). Some patterns of gender-specific enrichment were also observed for WHR. GI is the dominant tissue for females (*p* = 5.26×10^−5^) but seems less important in male samples (*p* = 2.95×10^−2^), while enrichment in muscle is consistent between males and females.

**Figure 4.**
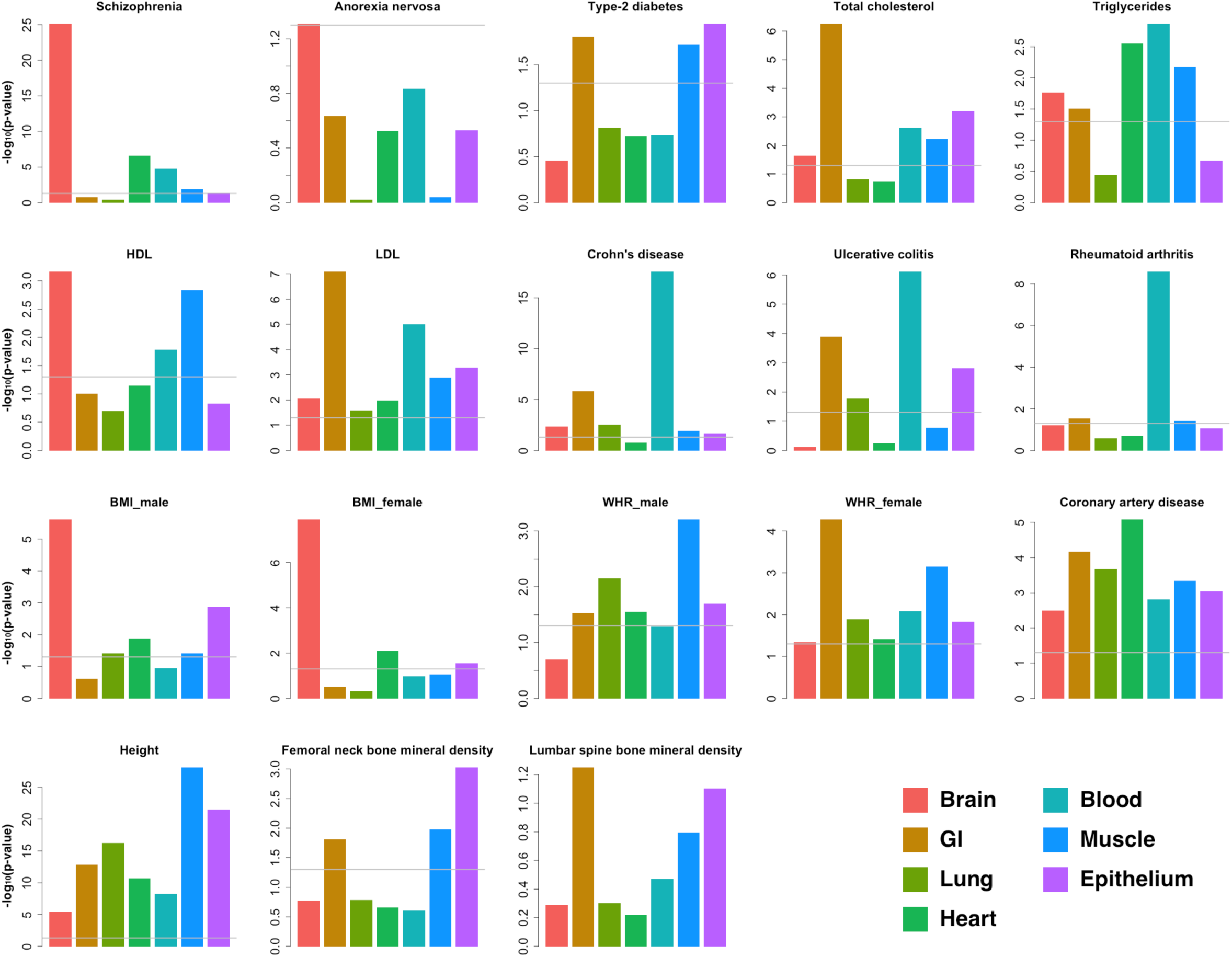
Tissue-specific enrichment of GWAS signals. Enrichment p-values were calculated using LD score regression. The grey line is the 0.05 cutoff for p-value.

It is worth noting that extra caution is needed when interpreting these enrichment results. For example, Finucane et al. reported connective/bone as the most enriched tissue type for human height^27^, but GenoSkyline annotations for this tissue is not available at this moment due to incomplete epigenomic data (**Online Methods**). Similarly, we are not yet able to investigate the relationship between lipid traits and liver tissue because of the lack of tissue-relevant functionality data.

### GWAS signal prioritization using tissue-specific functional annotations

We recently developed Genome Wide Association Prioritizer (GenoWAP), and showed that GWAS signals could be better prioritized through integrating GWAS summary statistics with GenoCanyon annotation^10^. From the results of tissue-specific enrichment analysis, it could be seen that some complex traits are strongly related to a few tissue types. In this section, we show that the performance of GWAS signal prioritization could be further improved through integrating GenoSkyline annotations of relevant tissue types.

Using both tissue-specific GS scores and GenoCanyon scores that quantify the overall functionality, we calculate the posterior probability *P*(*Z_D_* = 1, *Z_T_* = 1|*p*) to measure the importance of each SNP. In this calculation, Z_D_ is the indicator of disease/trait-specific functionality, Z_T_ is the indicator of tissue-specific functionality, and p is the p-value acquired from standard GWAS analysis (**Online Methods**). Psychiatric Genomics Consortium (PGC) has published two large GWAS meta-analyses for schizophrenia, a major psychiatric disorder. We applied our method to the smaller study^30^ and attempted to replicate the findings of the larger study^31^. This analysis demonstrates GenoSkyline’s ability to prioritize association signals that are more likely to be replicated in a larger sample. These two studies will be referred to as PGC2011 and PGC2014 studies in the following discussion.

Enrichment analysis suggests that brain is the most enriched of schizophrenia GWAS signals compared with other tissue types, and strong enrichment could also be observed in heart and blood (**Figure 4**). For each SNP in the PGC2011 study, mean GenoCanyon score of its surrounding region and mean GS scores of brain, blood, and heart tissues were calculated (**Online Methods**). SNPs in these tissue-specific functional regions and the SNPs in general functional regions are all enriched for associations with schizophrenia (**Figure 5a**; **Supplementary Figure 4**). Notably, tissue-specific functional regions are more enriched for associations with schizophrenia relative to general functional regions, with blood showing the strongest enrichment. It is also worth noting that non-functional regions are enriched of GWAS associations as well, most likely due to the LD between functional and non-functional SNPs^10^.

**Figure 5.**
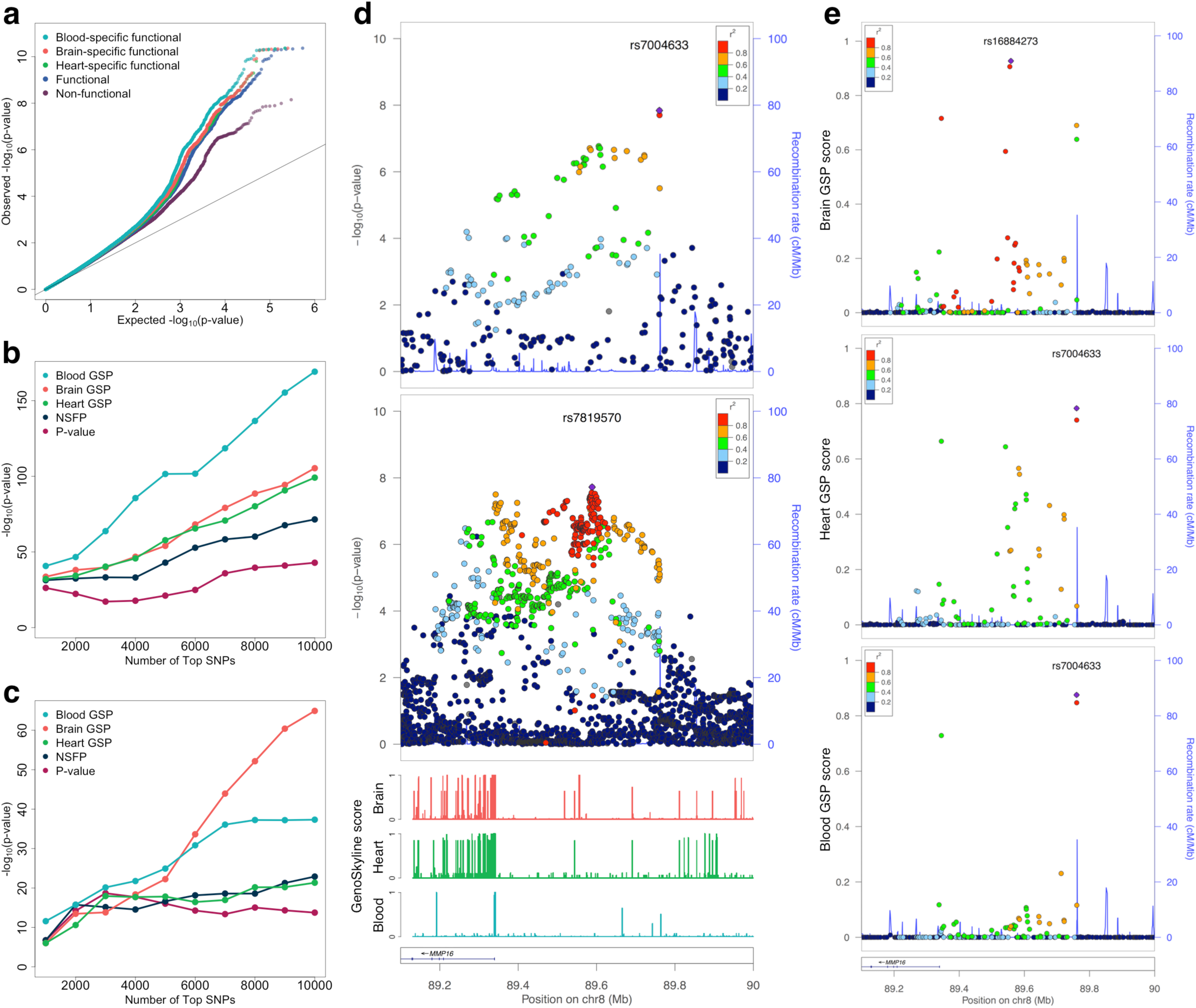
Prioritizing schizophrenia GWAS signals using GenoSkyline annotations. (a) Tissue-specific functional regions are more enriched of schizophrenia associations than generally functional regions and non-functional regions. (b) Enrichment of GTEx whole-blood eQTLs in top SNPs from PGC2011 study. (c) Enrichment of human brain quantitative trait loci in top SNPs from PGC2011 study. (d) Summary statistics at the schizophrenia-associated locus on chromosome 8q21 near *MMP16* gene. The top and middle panel show p-values from PGC2011 and PGC2-14 studies, respectively. The bottom panel shows GenoSkyline annotations at this locus. (e) Locus plots for tissue-specific posterior scores. From top to bottom, the three panels show posterior scores of brain, heart, and blood tissues, respectively.

Next, we define a new SNP-level metric for the tissue-specific GenoSkyline posterior (GSP) scores (i.e. *P*(*Z_D_* = 1, *Z_T_* = 1|*p*)) of brain, blood, and heart, as well as the nonspecific functionality posterior (NSFP) scores (i.e. *P*(*Z_D_* = 1|*p*); see **Online Methods**) for each SNP in PGC2011 study. Enrichment analysis using GTEx whole-blood eQTLs^11^ found that the top SNPs based on tissue-specific GSP scores are substantially more enriched of eQTLs than NSFP scores and p-values. As expected, blood GSP scores showed the strongest enrichment of whole-blood eQTLs (**Figure 5b**). When using a set of quantitative trait loci in human brain^32^, tissue-specific GSP scores also showed superior performance, with the brain-specific scores dominating others as the number of top SNPs increase (**Figure 5c** and **Supplementary Figure 5**).

A total of 108 schizophrenia-associated loci were identified in the PGC2014 study. We removed three loci on chromosome X due to the absence of SNPs on sex chromosomes in the PGC2011 dataset. All the SNPs in the PGC2011 study were ranked based on their p-values, NSFP scores, and tissue-specific GSP scores, respectively (**Supplementary Table 7**). The maximum ranks at each of the 105 schizophrenia-associated loci based on these different criteria were then compared (**Supplementary Table 8**). Brain GSP score showed better performance in prioritizing these loci when compared with p-value. Sixty-seven out of 105 loci had an increased rank (p-value=0.003, one-sided binomial test). The performance of heart GSP score was slightly worse than brain-specific score, but still better than p-value ranking. Blood GSP score showed comparable performance with p-value ranking. Notably, the performance of brain and heart GSP scores was still significantly better than NSFP score, although NSFP score outperforms ranking based on p-value.

Tissue-specific functional annotations could provide even deeper insight when prioritizing SNPs locally at risk loci. The schizophrenia-associated locus on chromosome 8q21 is located in the intergenic region upstream of *MMP16* gene (**Figure 5d**). The p-values in the PGC2014 study clearly suggested two signal peaks. One is located near the transcription start site of *MMP16,* while the other resides nearly 200,000 bases upstream and shows slightly stronger signal. However, the two-peak pattern was not clear in the PGC2011 study. Instead, two SNPs close to the end of the LD block near 89.8M showed the strongest signal. We compared the local predictions based on brain, heart, and blood-specific GSP scores at this locus (**Figure 5e**). Brain GSP scores successfully revealed the multi-peak nature at this locus, suggested the importance of the peak near 89.6Mb, and diminished the signal strength at the two SNPs near 89.8Mb, concurrent with the PGC2014 results. Although the method was applied on PGC2011 p-values, the results after prioritization matched the signal pattern in the PGC2014 study very well. Heart GSP scores also suggested the existence of the signal peak near 89.6M. However, the posterior scores have lower values, and the overall signal pattern does not match the PGC2014 study very well. The signal peak near 89.6M was completely lost in the blood-specific results. The two SNPs near 89.8M, however, had large GSP scores. The differences across tissue types are concurrent with GS scores at this locus (**Figure 5d**). Upstream of *MMP16,* near 89.6M, several functional segments can be seen in brain, only one remains in heart, and none exists in blood. Through comparing the tissue-specific prioritization results with the p-values in PGC2014 study, we see that brain-specific GSP scores had the strongest signal strength, which can be quantified using the local maximum GSP score (**Online Methods**). The highly matched signal pattern also suggested that brain might be the tissue type in which this locus plays a functional role.

### Further insight on risk loci associated with coronary artery disease

Next, we applied our method to another GWAS to further illustrate the biological insight that GenoSkyline can provide for understanding complex diseases. The CARDIoGRAM consortium published a large-scale GWAS meta-analysis of coronary artery disease (CAD) comprising 22,333 cases and 64,762 controls^33^, in which they replicated 10 out of 12 previously reported risk loci and identified 13 new loci associated with CAD. We applied our method on the summary statistics and used the local maximum GSP score to measure the relatedness between each risk locus and different tissue types (**Online Methods**). We removed the locus on chromosome 1q41 (*MIA3*) and the locus on chromosome 6q25.3 (*LPA*) due to incomplete data in the meta-analysis stage of CARDIoGRAM study. The remaining 21 CAD-associated loci are summarized in **Table 1** and **Supplementary Table 9**.

**Table 1.**
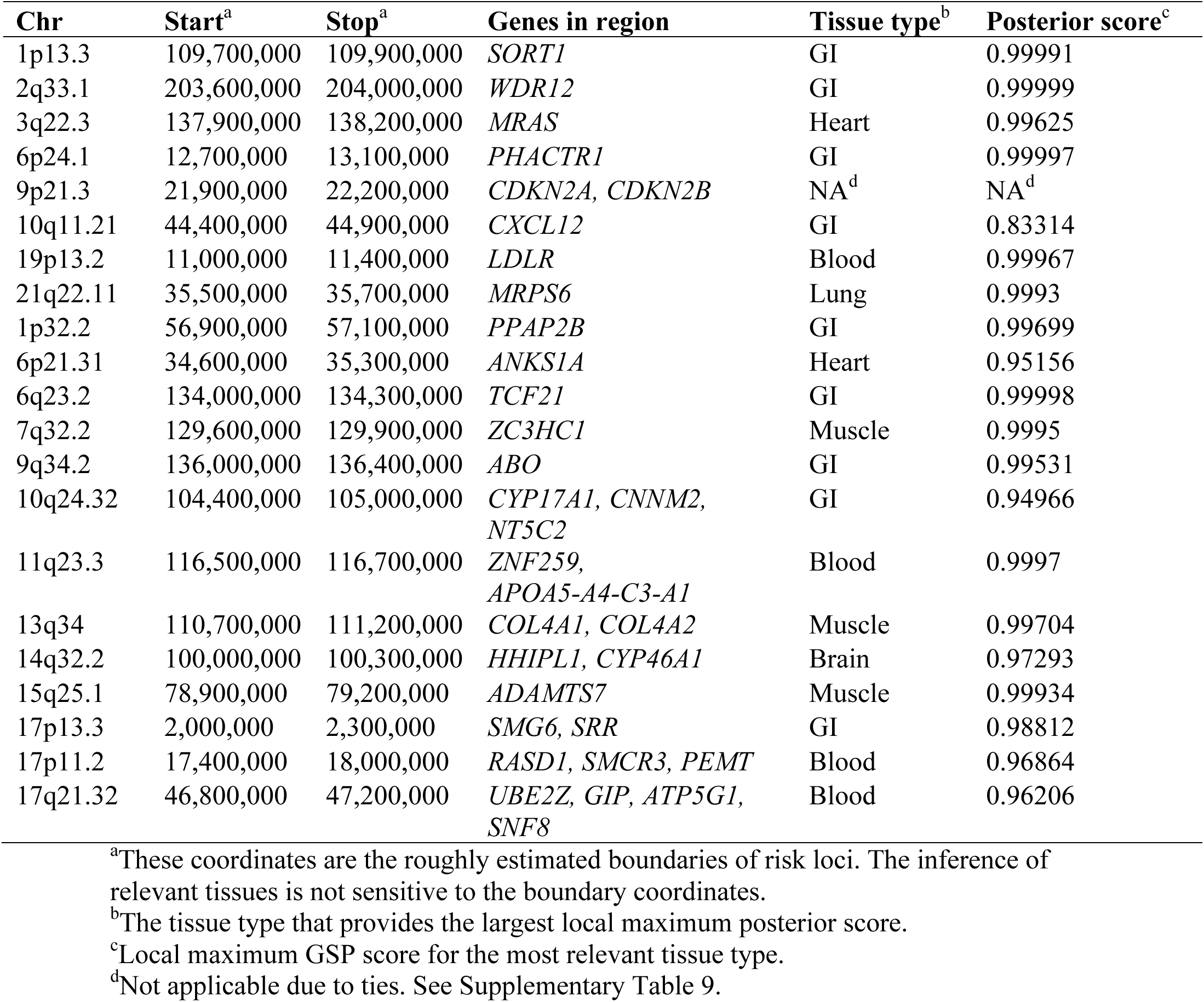
Risk loci for coronary artery disease

The first impression of these results is that despite the strong overall enrichment of GWAS signals (**Figure 4**), heart is the most relevant tissue type for only two loci. On the contrary, a substantial proportion of risk loci (9 out of 21) seem to be functional in the GI tissue. Interestingly, GI was the most enriched tissue type for several known risk factors for CAD including LDL and total cholesterol (**Figure 4**). These results suggest not only the larger effect sizes of CAD-associated loci in the gastrointestinal system, but also a substantial amount of undetected heart-related loci. Furthermore, brain was the least enriched tissue type for CAD GWAS signals, but the risk locus on chromosome 14q32.2 near *HHIPL1* and *CYP46A1* was predicted to be functional in brain. In fact, the *CYP46A1* gene encodes for Cholesterol 24-hydroxylase that is present mainly in brain, where it converts cholesterol from degraded neurons into 24S-hydroxychoelesterol^33,34^. This process is crucial for eliminating cholesterol from the brain since cholesterol is usually unable to pass the blood-brain barrier^35^.

A larger GWAS for CAD was published during the preparation of this manuscript^36^. This large study may be used to validate the performance of our approach when its summary statistics become publicly available in the future.

## Discussion

In this paper, we introduced GenoSkyline, an integrative framework for predicting tissue-specific functional regions in the human genome. Through integrating GenoSkyline annotations with GWAS summary statistics, we illustrated a variety of ways that GenoSkyline could help researchers understand human complex diseases and traits. We also showed that the GenoSkyline framework is customizable so that researchers can develop their own functional annotations for a selected group of cells. As epigenomic ChIP-seq data become available for an increasing number of cell types in the future, GenoSkyline’s ability to facilitate studies of complex disease will be further enhanced.

Our approach is not without limitation. First, the annotation results are incomplete due to currently unavailable tissue types, and as a result, the GWAS enrichment results may not be comprehensive (e.g. liver may also be highly related to CAD, but there is no complete annotation data from liver yet). Second, some risk loci (or independent functional segments at the same locus) may play active roles in multiple tissue types. For example, in our PGC GWAS analysis, although local maximum GSP scores suggest that brain may be more relevant with the risk locus upstream of *MMP16,* two SNPs near 89.8MB are located near several functional segments in blood. Whether these SNPs can be functionally linked to schizophrenia remains to be investigated. Moreover, we emphasize that our method identifies regions of likely functionality, but does not provide conclusive proof of functionality for any individual SNP or locus. That said, our method still provides a simple and intuitive summary statistic that measures the relatedness between risk loci and sets of functionally related tissues. It has great potential to become a standard step in downstream GWAS analysis to help researchers generate new hypotheses regarding the etiology behind each risk locus.

The increasing accessibility of GWAS summary statistic datasets, coupled with the method’s independence from requiring individual-level genotype and phenotype, make Genoskyline tissue-specific prioritization useful and easy to implement. Moreover, GWAS signal integration is just one way to utilize GenoSkyline annotations. Its nucleotide-level functional prediction based on unsupervised learning and the good predictive performance in non-coding regions promise a potential role in many fields of genomics, such as next-generation sequencing studies and understanding somatic mutations. GenoSkyline scores of seven tissue types and two additional cell types have been pre-calculated for the entire human genome and can be readily downloaded. Source code is available for all major OSes and can be accessed at (http://genocanyon.med.yale.edu/GenoSkyline). We believe that GenoSkyline and its applications can guide genetics research at multiple resolutions and greatly benefit the broader scientific community.

## Online Methods

### Consolidated epigenomes

Epigenetic data were selected from the Epigenomics Roadmap Project’s 111 consolidated reference epigenomes database^3^ (http://egg2.wustl.edu/roadmap/) based on anatomy type and mark availability. Each tissue type is a clustering of relevant samples in order to contain at least one of each of the following: H3k4me1, H3k4me3, H3k36me3, H3k27me3, H3k9me3, H3k27ac, H3k9ac, and DNase I Hypersensitivity. Samples are reduced to a per-nucleotide binary encoding of presence or absence of narrow contiguous regions of ChIP-seq signal enrichment compared to input (Poisson p-value threshold of 0.01), and a union of all tissue-specific samples for that mark is taken. The set of 8 marks was chosen due to the well-understood, localized regulatory interactions of histone marks^37^ and DNase I^38^. We created nine unique tissue and cell type clusters based on these annotations (**Supplementary Table 1**) to represent common, physiologically-related organ systems. To reflect actual tissue-specific epigenetic behavior, a majority of samples chosen are primary tissues and cultures, and inclusion of immortalized cell lines has been kept to a minimum.

### GenoSkyline model and estimation

Lu et al. previously proposed a method that applies unsupervised-learning techniques on genomic annotations to predict the functional potential of a genomic region^4^. Given a set of annotations ***A***, we assume the joint distribution of ***A*** along the genome to be a mixture of annotations at locations with no functionality, i.e. *f*(***A*** | *Z* = 0), and annotations at locations that are functional, i.e. *f*(***A*** | *Z* = 1). We assume that each annotation in ***A*** is conditionally independent given Z, allowing the conditional joint density of ***A*** given Z to be factorized as

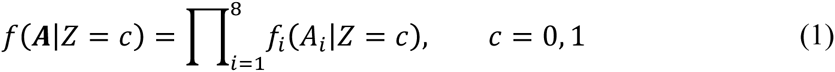

Since all annotations used are binary classifiers, the Bernoulli distribution was used to model the marginal functional likelihood given each individual annotation.

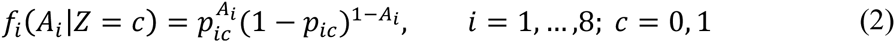

Assuming a prior probability *π* of being functional (*π* = *P*(*Z* = 1)), we can estimate the parameter *p_ic_* of each annotation with the Expectation-Maximization (EM) algorithm, and calculate the posterior probability at a given genomic coordinate, referred to as the GS score.

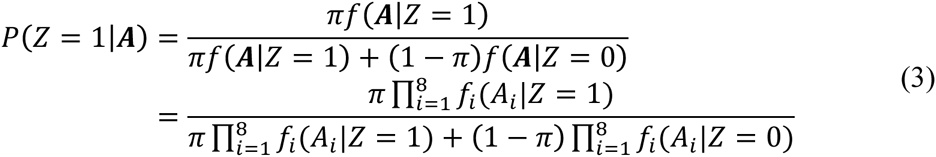

We must estimate 17 parameters for each tissue tract.

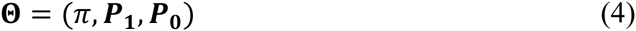

Where

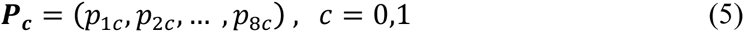

Parameters were estimated using the GWAS Catalog^6^, downloaded from the NHGRI website (http://www.genome.gov/gwastudies/), which at the time of download, contained 13,070 unique SNPs found to be significant in at least one published GWAS. These SNPs were expanded into 1k bp intervals, and formed a genome sampling covering 12,801,840 bp of the genome. While significant SNP associations are likely to tag the effects of nearby functional elements, the size and distance of these functional elements varies for each individual SNP. As a result, the total sampling serves as an effective and robust representation of functional and non-functional regions along the genome (**Supplemental Notes**).

### Case studies of experimentally validated functional machinery

VISTA enhancers^14^ were downloaded from the VISTA Enhancer Browser (http://enhancer.lbl.gov/), where enhancers with E11.5 reporter staining experimental data were selected. Brain enhancers were selected based on staining results identifying any CNS-related tissues (neural tube, cranial nerve, hindbrain, mesenchyme derived from neural crest, trigeminal V, forebrain, and midbrain). Heart enhancers were enhancers identified for positive reporter results in the heart region of E11.5 mouse reporter assays. Blood vessels enhancers were identified by selecting for “blood vessels” expression pattern. Hypomethylated TE loci in H1ES, brain, breast, and blood were downloaded from http://epigenome.wustl.edu/TE_Methylation/. All genomic coordinates were converted to genome build hg19.

### SNP prioritization using tissue-specific functional annotation

We identify three disjoint cases for a given GWAS SNP:

1. The SNP is in a genomic region that is functional for the given phenotype and tissue (Z_D_ = 1, Z_T_ = 1).
2. The SNP is in a genomic region that is functional for the given tissue, but that tissue has no functionality in the phenotype (Z_D_ = 0, Z_T_ = 1).
3. The SNP is in a genomic region that is not functional in the given tissue (Z_T_ = 0).

A useful metric for prioritizing SNPs is the conditional probability that the SNP is classified under case-I given its p-value in a given GWAS study, i.e. *P(Z_D_ = 1, Z_T_ = 1|p*). We can calculate this probability by employing Bayes formula and considering all three cases as follows:

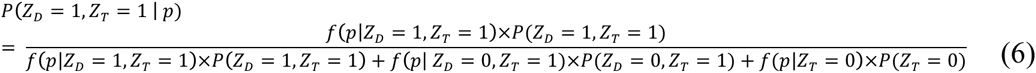

First, the case in which Z_T_ = 0 can be directly identified by assigning each SNP a prior probability of tissue-specific functionality (i.e. *P(Z_T_ =* 1)) defined as the average GS score of its surrounding 10,000 base pairs for that tissue (**Supplementary Notes**). We partition all the SNPs into two subgroups based on a mean GS score threshold of 0.1, although these probabilities take on a bimodal distribution and are not sensitive to changing threshold^10^. In this way, we can use these partitions to directly estimate f(p|Z_T_ = 0) by applying density estimation techniques on the SNP subgroup with low GS scores. More specifically, we apply histogram for density estimation and use cross validation to choose the optimal number of bins.

Second, we estimate the p-value density of our second case, where Z_D_ = 0 and Z_T_ = 1. We can intuitively assume that SNPs that are functional in a tissue but not relevant to the phenotype will have similar p-value behavior to all other SNPs that are not relevant to the phenotype, which in turn behave similarly to SNPs that are not functional at all (**Supplementary Notes**). More formally, we can describe this relationship as follows:

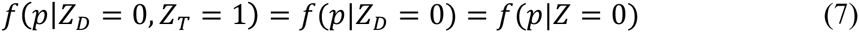

We can effectively estimate *f*(*p|Z* = 0) by using a similar approach to estimating *f*(*p|Z_T_* = 0), but partitioning SNPs using the general functionality GenoCanyon score instead of tissue-specific GS score.

Next, we consider the following formulas.

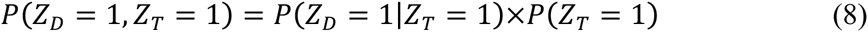

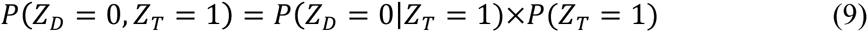

The prior probability *P*(*Z_T_* = 1) can be calculated directly from GS scores as stated above, but the conditional probabilities of disease-specific functionality given tissue-specific functionality remains to be estimated.

Finally, we estimate all the remaining terms in formula 4 using the EM algorithm. In the first step of the estimation procedure, we acquired the subset of SNPs located in tissue-specific functional regions. The p-value distribution of these SNPs is the following mixture.

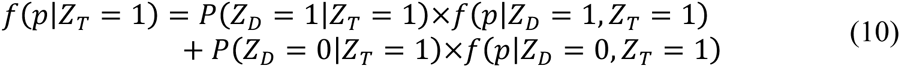

Density *f*(*p|Z_D_* = 0, *Z_T_* = 1) has been estimated in earlier steps. Applying the findings of Chung et al., we assume a beta distribution of the p-values of functional SNPs (i.e. *f*(*p|Z_D_* = 1, *Z_T_* = 1)) as a reasonable approximation under general assumptions of SNP effect size^9^.

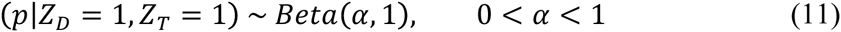

The EM algorithm is then applied to the SNP subset located in tissue-specific functional regions. The beta assumption guarantees a closed-form expression in each iteration and all the remaining parameters can be subsequently estimated. We now have all the necessary terms for equation 4, and define this as our posterior probability score of tissue-specific disease functionality (GSP score). The feature of integrating tissue-specific functional annotations to prioritize GWAS signals has been added to the GenoWAP software available on our server (http://genocanyon.med.yale.edu/GenoSkyline).

### SNP prioritization using GenoCanyon annotation

Non-tissue specific GenoCanyon scores are assigned to GWAS signals using GenoWAP^10^. Briefly, GenoWAP calculates the posterior score *P*(*Z_D_* = 1|*p*) using a simpler model for functionality.

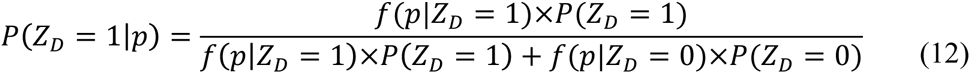

This conditional probability can be calculated similarly to GS scores, making use of equation (5) to empirically estimate *f*(*p*|*Z_D_* = 0), a beta distribution on partitioned Genocanyon scores (calculated with 22 tissue non-specific ENCODE functionality annotations^4^) to estimate *f*(*p*|*Z_D_* = 1), and the EM algorithm on the functional marker p-value density to calculate *P*(*Z_D_* = 1) as described in Lu *et al.* These are referred to in the results as the NSFP scores to which GenoSkyline SNP prioritization is compared.

### Calculating tissue-specific enrichment using LD score regression

Enrichment of GenoSkyline-derived tissue-specific annotations in GWAS summary statistics was calculated using stratified LD score regression^27^. First, tissue-specific annotations were computed using GenoSkyline scores, 1000 Genomes data of European ancestry^39^ and a 1-centiMorgan (cM) window. Then the annotations were analyzed by adding each one of them to the full baseline model to control for 53 categories of general annotations. For each tissue-specific annotation, partitioned heritability was estimated using stratified LD score regression^27^ and enrichment was then calculated as the ratio of proportion of SNP heritability explained by the annotation and proportion of SNPs in the annotation.

### Measuring relevant tissue types for GWAS risk loci

A large GSP score is obtained if the p-value for the SNP is small and the SNP is located in a highly functional region for the tissue type under investigation. Therefore, the maximal GSP score at a risk locus effectively measures how well the p-values match the pattern of GenoSkyline annotations, thereby measuring the relatedness between the GWAS locus and different tissue types. For each tissue, the maximal GSP score is acquired at the risk locus of interest. These scores are then compared across tissue types. The largest score is referred to as local maximum GSP score, and the corresponding tissue type is predicted to be the most relevant tissue.

### Bioinformatics tools

Locus plots were generated using LocusZoom^40^. The “ggbio” R package^41^ was used to plot genes. The “bigmemory” R package^42^ was used to access and manipulate massive dataset.

## Acknowledgements

This study was supported in part by the National Institutes of Health grants R01 GM59507, the VA Cooperative Studies Program of the Department of Veterans Affairs, Office of Research and Development, and the Yale World Scholars Program sponsored by the China Scholarship Council.

## Author Contributions

Q.L. and R.L.P. conceived the project, wrote the initial draft, and performed the analyses. Q.W. performed tissue-specific enrichment analysis. B.J.H. performed one heart-related case study. H.Z. advised on statistical and genetic issues.

## Supplementary Notes

### Bimodal pattern in GenoSkyline score

When estimating the proportion of functional genome for each tissue type, we adopted 0.5 as the cutoff for GS score. The GS score histograms for different tissues on chromosome 22 are plotted in **Supplementary Figure 6**. The GS score distributions have a clear and consistent bimodal pattern across different tissue types. The similar bimodal pattern can also be observed for other chromosomes. Therefore, the cutoff choice does not substantially affect the estimation of functional proportion.

### Robustness of GenoSkyline parameter estimation

The parameters in the GenoSkyline framework were estimated from a set of 12,801,840 bases acquired from GWAS catalog. The reason of using this GWAS-based set is to guarantee the inclusion of a sufficient amount of functional bases. In our previous work, we showed that parameter estimation under this framework is robust^4^. Here, all the 17 parameters were re-estimated after adding 2,000,000 and 6,000,000 bases randomly selected from chromosome 1 to the initial set containing 12,801,840 bases. The parameter estimates remained highly stable (**Supplementary Table 10**). These results show that GenoSkyline parameter estimation is insensitive to the choice of the initial set.

### Several remarks on GSP score

For GSP score calculation, we used the mean GS score of the surrounding 10,000 bases as the prior probability *P*(*Z_T_* = 1) for each SNP. This is because the nucleotide-level GS score may be insufficient for GWAS signal prioritization. In fact, each SNP in GWAS carries information of its nearby variants that are not genotyped or imputed. A better-informed metric needs to measure the functional potential for the surrounding region of each SNP. We chose 10,000 bases as the window size, but no substantial difference in empirical performance was observed when changing the window size to 5,000 or 20,000. In our implemented GenoWAP software for SNP prioritization (available at http://genocanyon.med.yale.edu/GenoSkyline), the users are allowed to use their own annotation data. Therefore, the window size can be changed when necessary. Since the mean GS score of surrounding regions was used as the prior, our SNP prioritization approach is in fact a region-based tool. We emphasize again that it identifies regions of likely functionality and substantially improves the resolution of GWAS, but does not provide conclusive proof of functionality for any individual SNP or locus.

In order to calculate the GSP score, we assumed that SNPs that are functional in a tissue not relevant to the phenotype would have similar p-value behavior to all other SNPs that are not relevant to the phenotype, which in turn behave similarly to SNPs that are not functional at all (see equation 7 in **Online Methods**). This assumption may not hold exactly due to some intrinsic differences between SNPs located in non-functional regions (Z=0) and those in non-specific functional (Z=1) and tissue-specific functional regions (Z_T_=1). As far as we are aware, the main possible contributing factor may be different linkage disequilibrium (LD) patterns in those regions with Z = 0 and those with Z = 1 or Z_T_=1. For example, if there is stronger LD in Z_T_ = 1 regions, then the markers with Z_D_ = 0 and ZT = 1 may have a different p-value distribution from those with Z = 0.

In order to check if this is a serious issue, we compared the LD patterns in regions with Z = 0, Z =1, and Z_T_ = 1 for multiple tissue types on chromosome 22. We downloaded the pre-calculated LD scores^43^ for the 1000 Genomes European population from the LD score GitHub page (https://github.com/bulik/ldsc/wiki/LD-Score-Estimation-Tutorial). Based on cutoff 0.1 for GenoCanyon and GenoSkyline scores, we divided all the SNPs on chromosome 22 into tissue-specific functional, non-specific functional and nonfunctional subcategories. The kernel density estimates of the two subgroups are plotted in **Supplementary Figure 7**. It can be seen that there is no substantial difference of LD score distributions between different categories. Therefore, it is reasonable to assume the LD patterns in regions with Z = 0, Z = 1, and Z_T_ = 1 to be similar. Moreover, as Z_D_ = 1 is a relatively small proportion of Z = 1, this assumption is likely to be a good approximation.

**Supplementary Figure 1.**
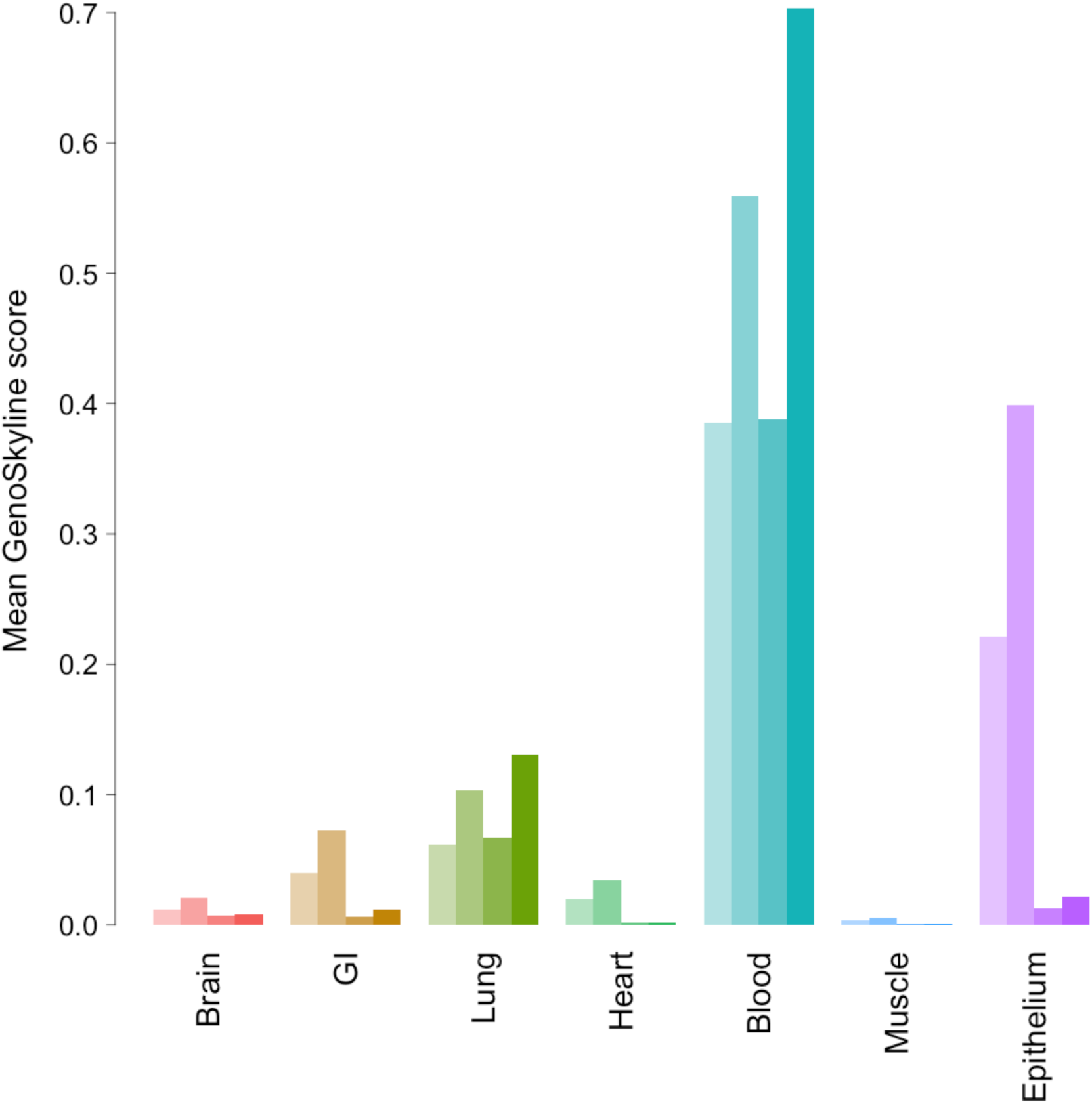
Mean GS score for different region categories in *HBB* gene complex across seven tissue types. For each tissue, the four bars from left to right indicate all 23 CRMs, adult CRMs, all genes, and adult globins, respectively.

**Supplementary Figure 2.**
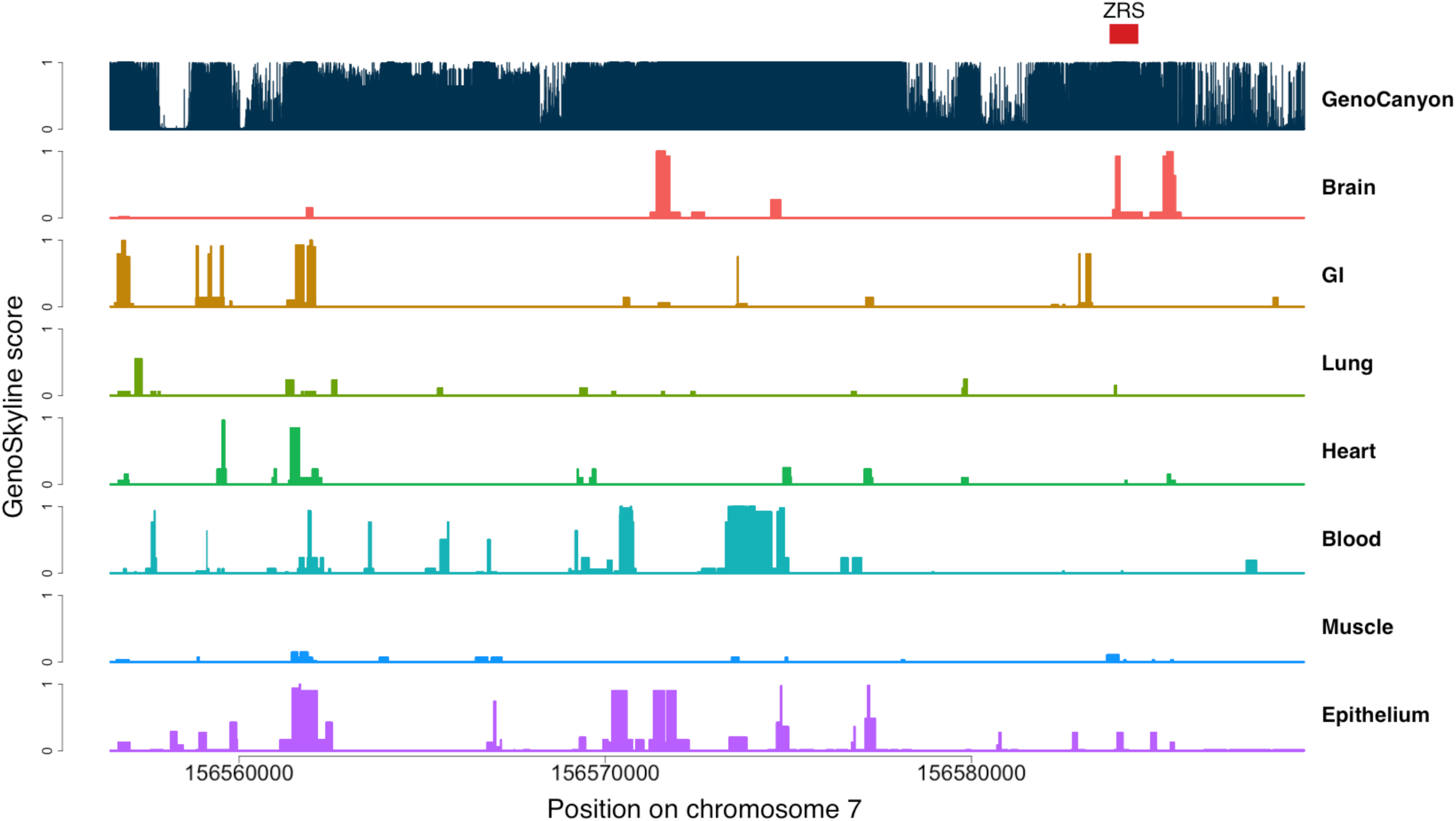
GenoCanyon score and GenoSkyline scores for seven tissues in the 5th intron of *LMBR1.* The red box marks the location of ZRS.

**Supplementary Figure 3.**
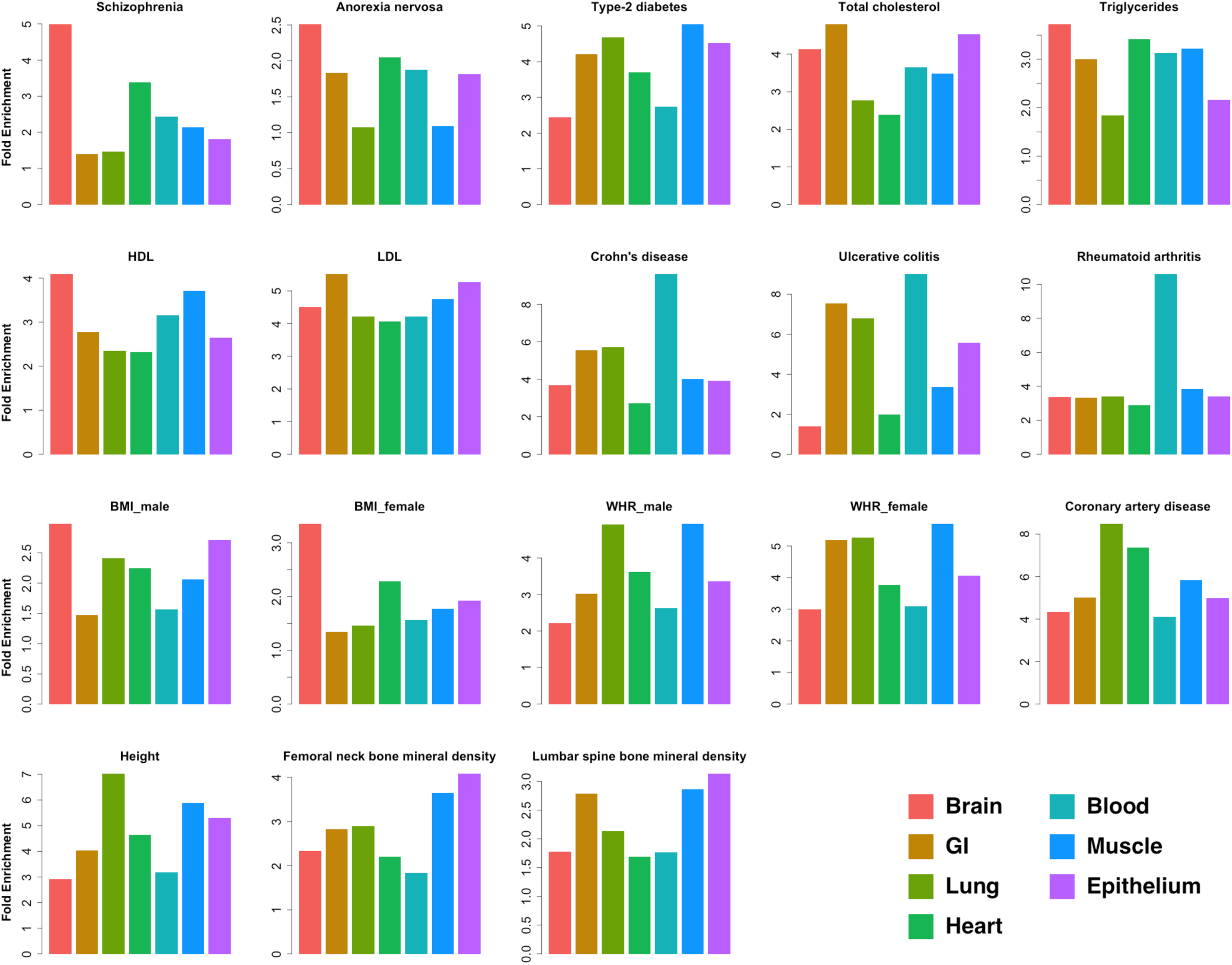
Tissue-specific fold enrichment of GWAS signals.

**Supplementary Figure 4.**
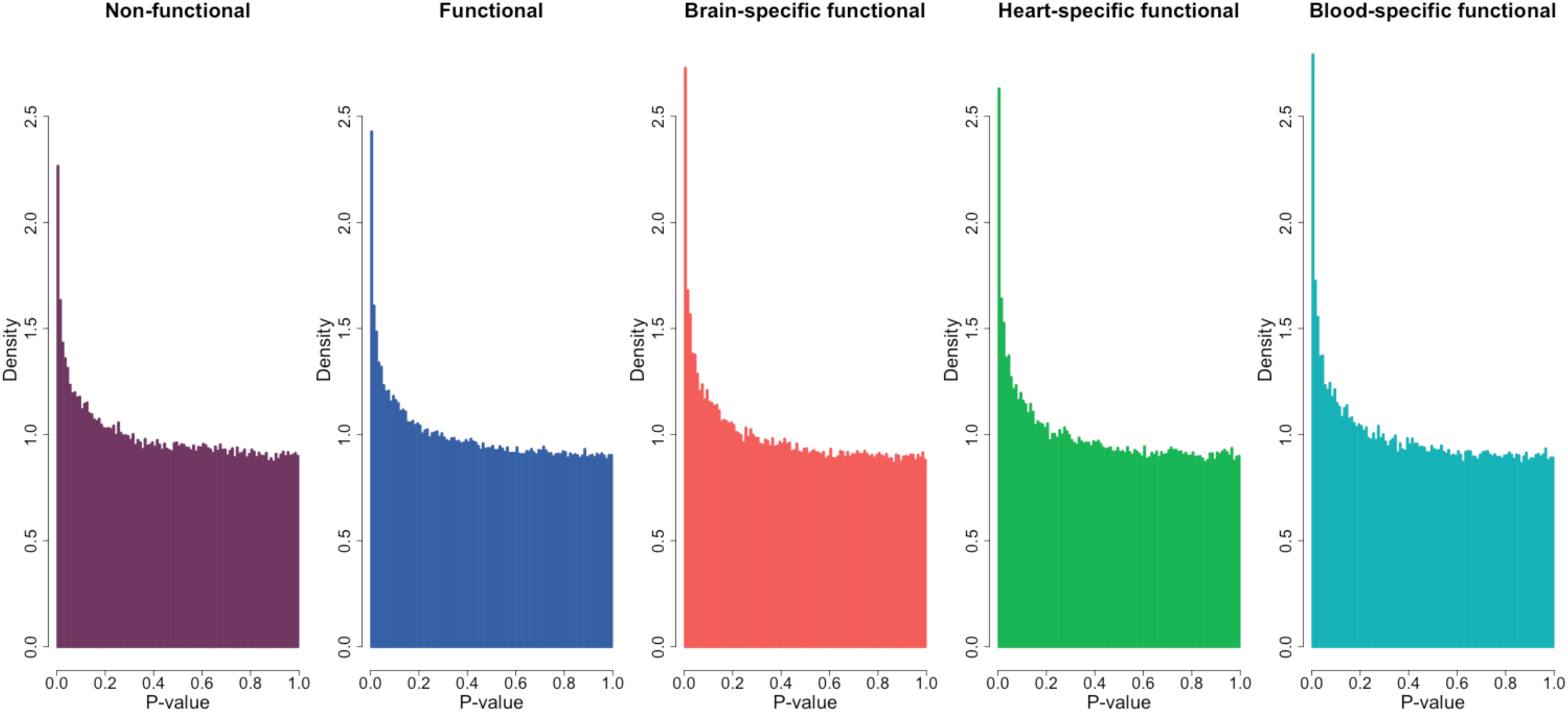
Histograms of p-values for SNPs located in non-functional, functional, and tissue-specific functional regions.

**Supplementary Figure 5.**
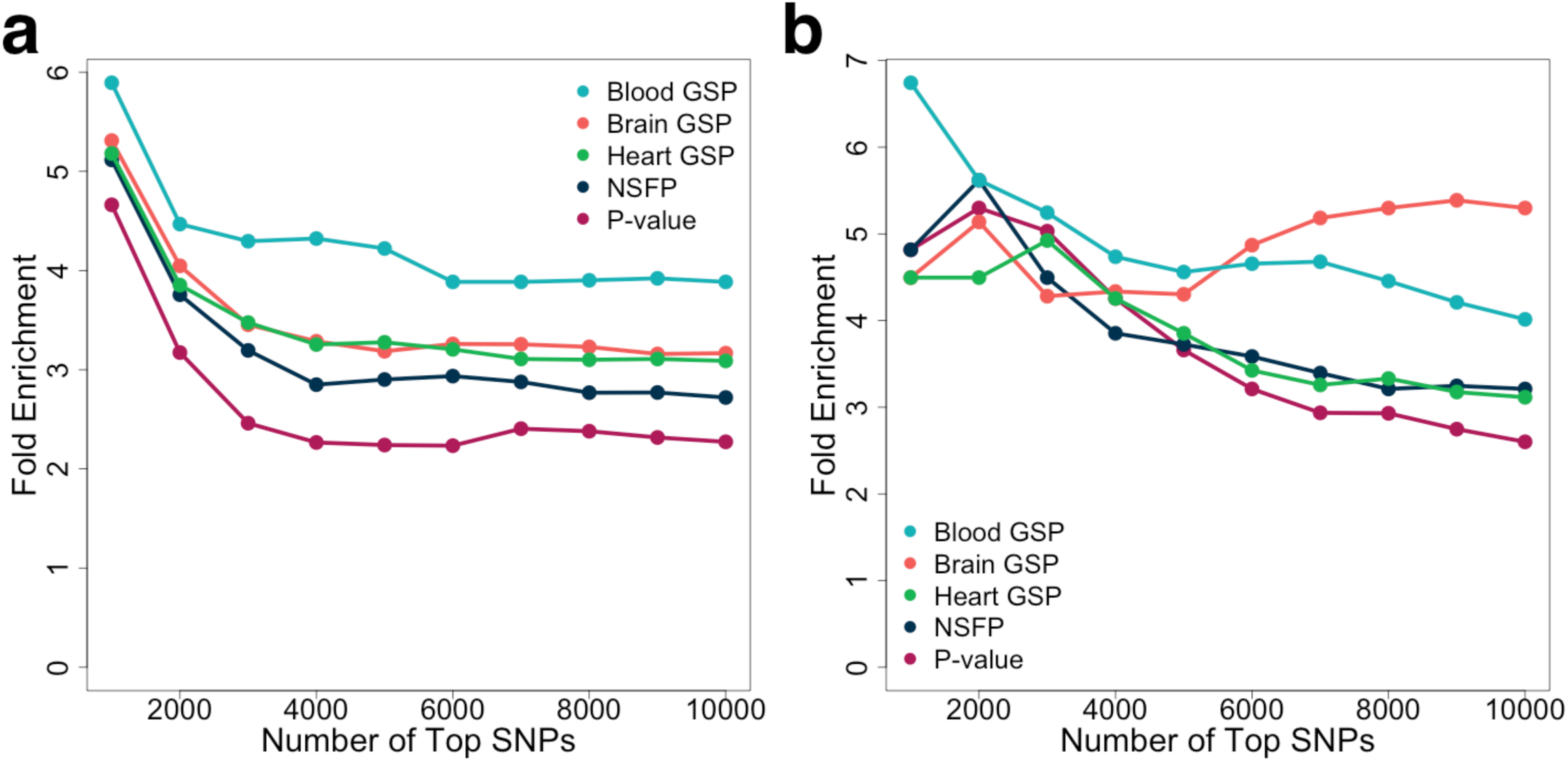
Fold enrichment of eQTLs in top SNPs from PGC2011 study. (a) GTEx whole-blood eQTLs. (b) Human brain quantitative trait loci.

**Supplementary Figure 6.**
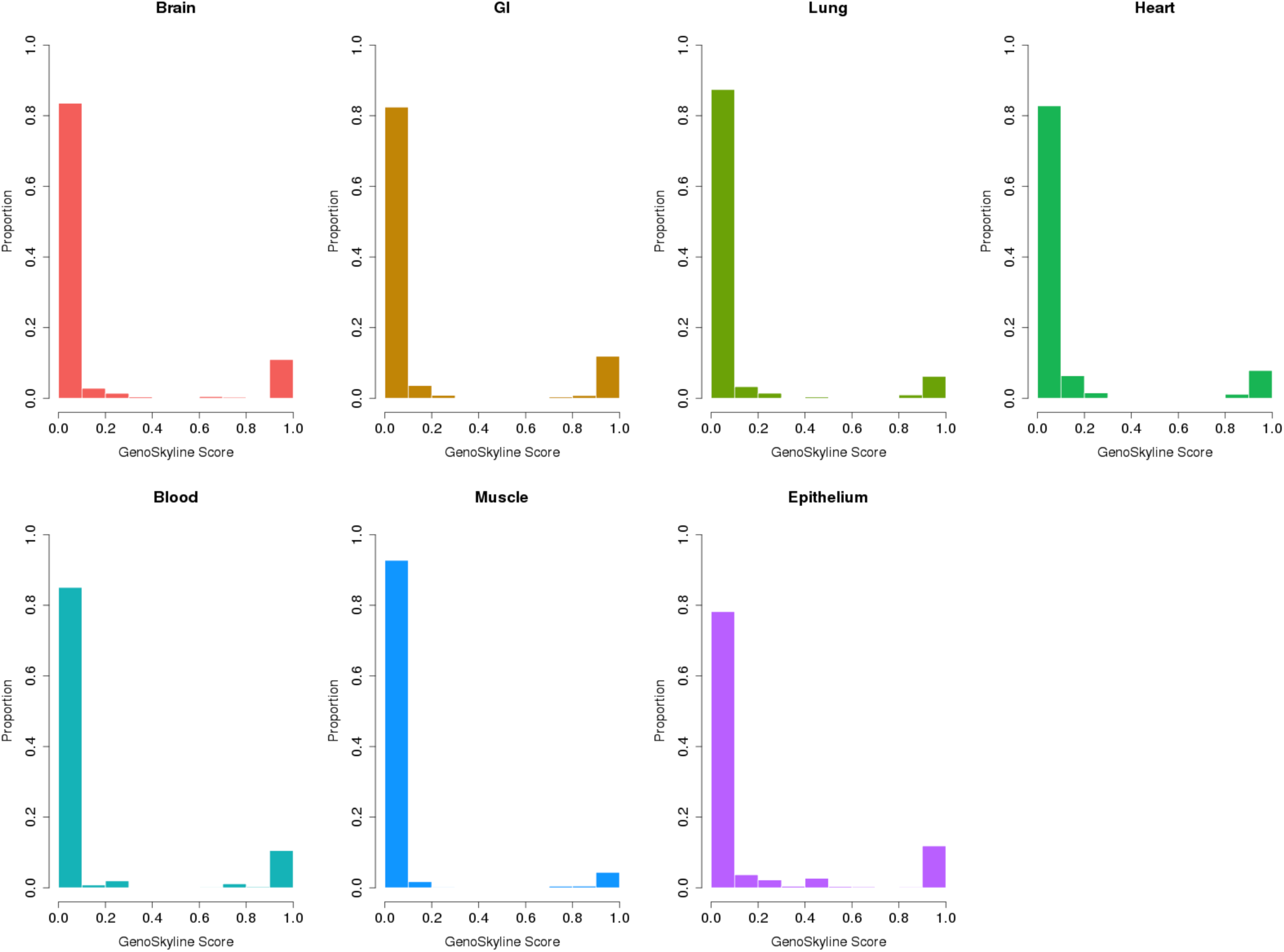
Distribution of GenoSkyline scores on chromosome 22.

**Supplementary Figure 7.**
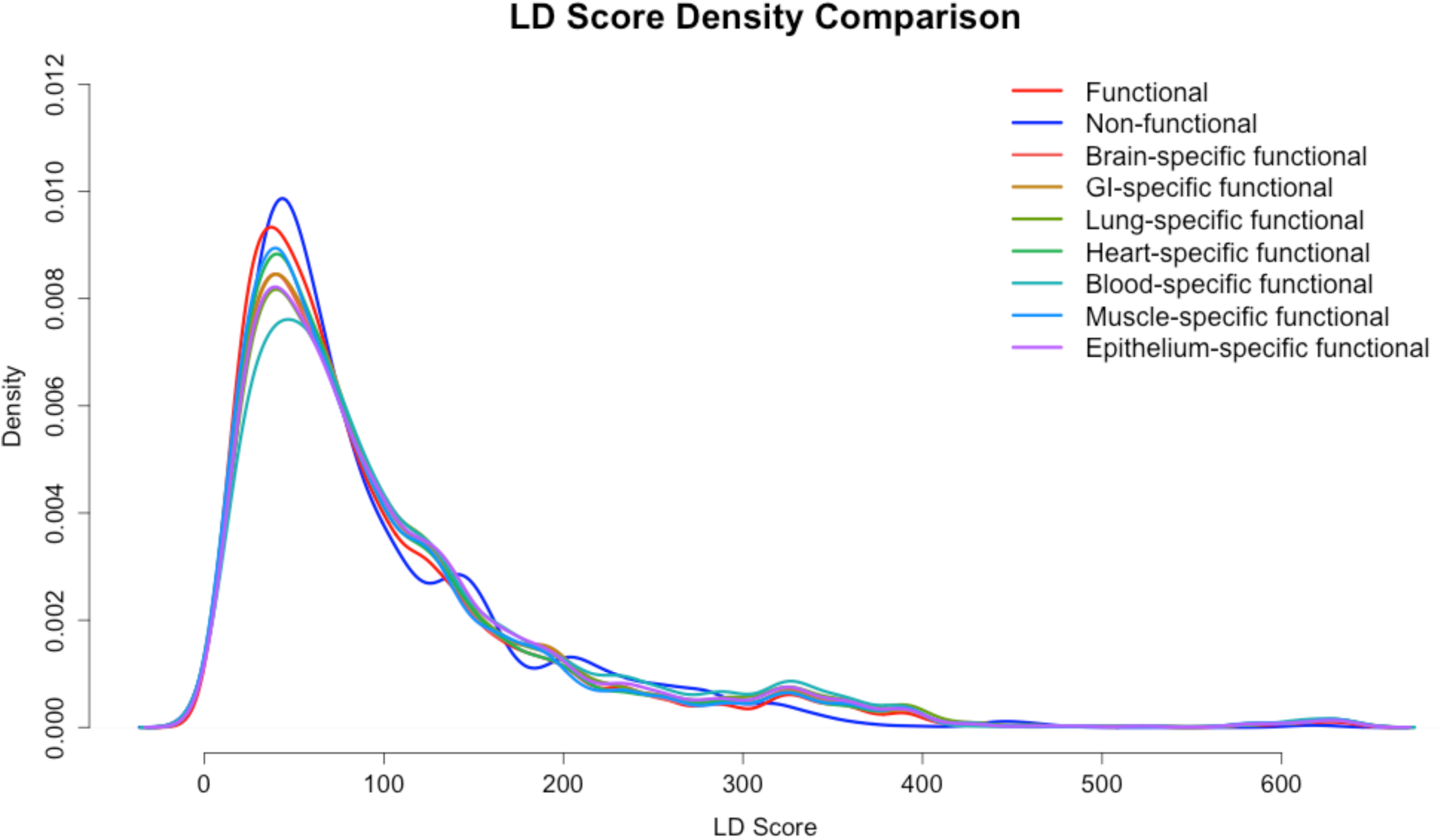
Comparison of LD score densities on chromosome 22 across different SNP categories.

**Supplementary Table 1.**
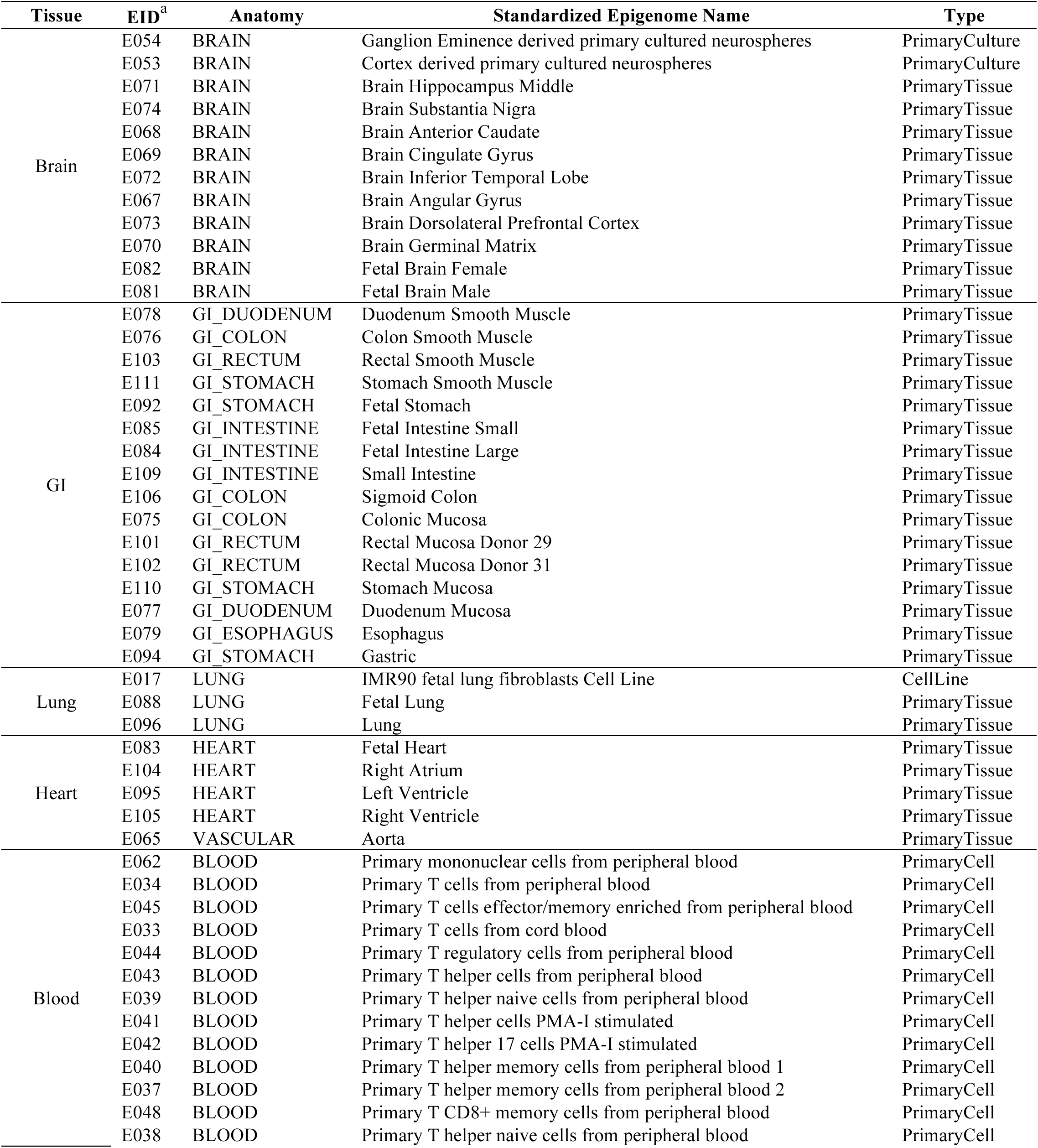
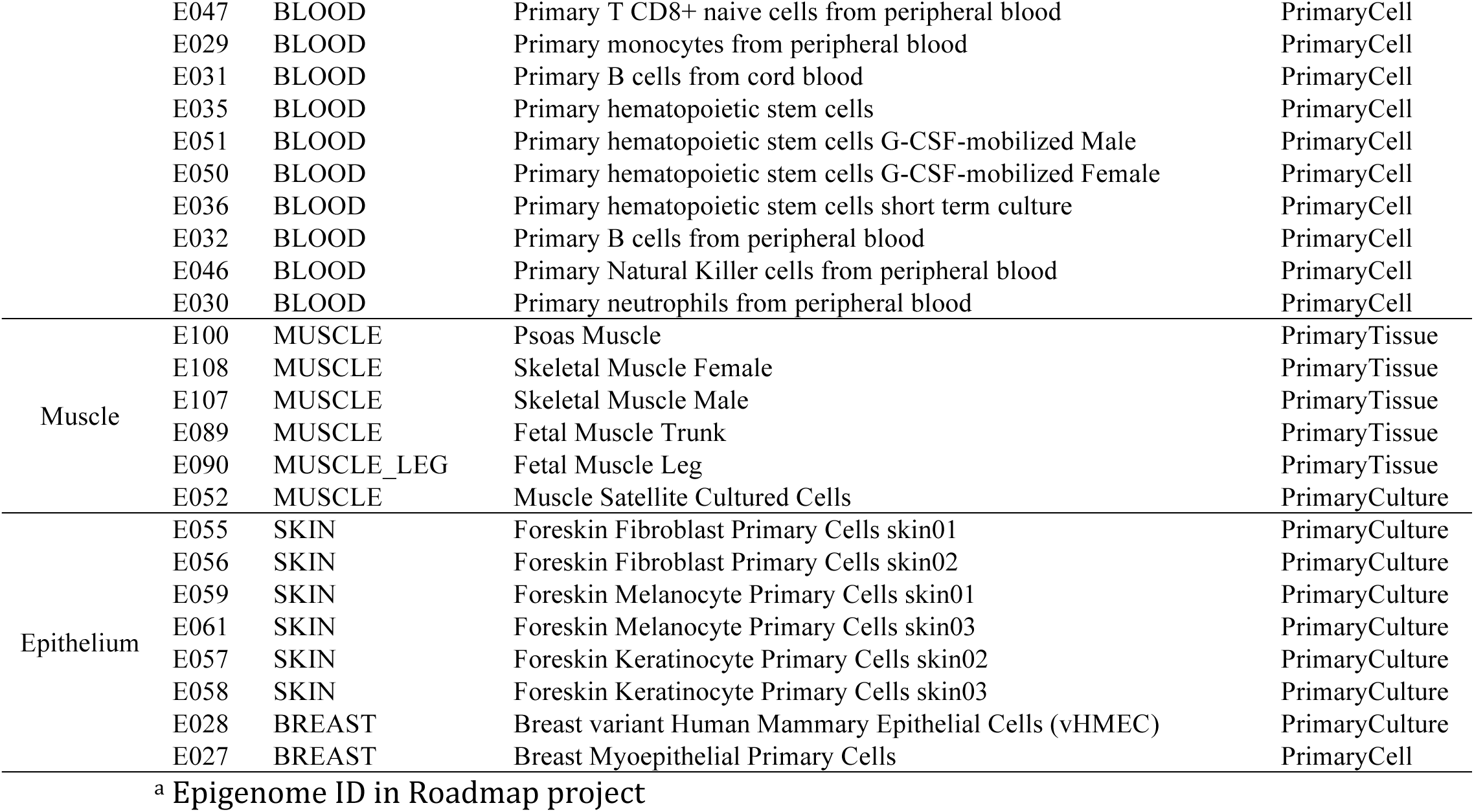
Cell types used for developing GenoSkyline annotations of seven tissue types.

**Supplementary Table 2.**
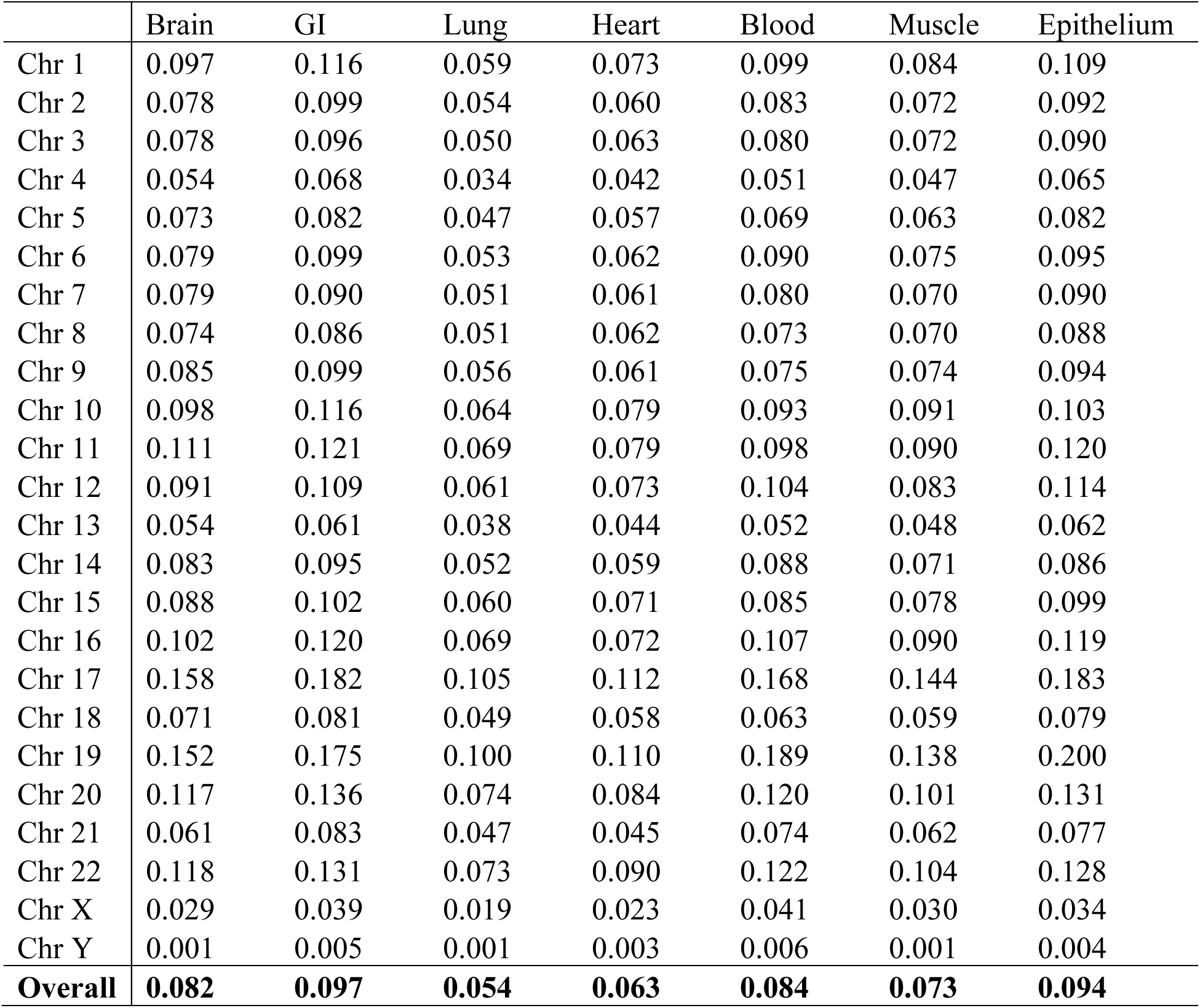
Proportion of functional genome across seven tissue types under GS score cutoff 0.5.

|  | Brain | GI | Lung | Heart | Blood | Muscle | Epithelium |
| --- | --- | --- | --- | --- | --- | --- | --- |
| Chr 1 | 0.097 | 0.116 | 0.059 | 0.073 | 0.099 | 0.084 | 0.109 |
| Chr 2 | 0.078 | 0.099 | 0.054 | 0.060 | 0.083 | 0.072 | 0.092 |
| Chr 3 | 0.078 | 0.096 | 0.050 | 0.063 | 0.080 | 0.072 | 0.090 |
| Chr 4 | 0.054 | 0.068 | 0.034 | 0.042 | 0.051 | 0.047 | 0.065 |
| Chr 5 | 0.073 | 0.082 | 0.047 | 0.057 | 0.069 | 0.063 | 0.082 |
| Chr 6 | 0.079 | 0.099 | 0.053 | 0.062 | 0.090 | 0.075 | 0.095 |
| Chr 7 | 0.079 | 0.090 | 0.051 | 0.061 | 0.080 | 0.070 | 0.090 |
| Chr 8 | 0.074 | 0.086 | 0.051 | 0.062 | 0.073 | 0.070 | 0.088 |
| Chr 9 | 0.085 | 0.099 | 0.056 | 0.061 | 0.075 | 0.074 | 0.094 |
| Chr 10 | 0.098 | 0.116 | 0.064 | 0.079 | 0.093 | 0.091 | 0.103 |
| Chr 11 | 0.111 | 0.121 | 0.069 | 0.079 | 0.098 | 0.090 | 0.120 |
| Chr 12 | 0.091 | 0.109 | 0.061 | 0.073 | 0.104 | 0.083 | 0.114 |
| Chr 13 | 0.054 | 0.061 | 0.038 | 0.044 | 0.052 | 0.048 | 0.062 |
| Chr 14 | 0.083 | 0.095 | 0.052 | 0.059 | 0.088 | 0.071 | 0.086 |
| Chr 15 | 0.088 | 0.102 | 0.060 | 0.071 | 0.085 | 0.078 | 0.099 |
| Chr 16 | 0.102 | 0.120 | 0.069 | 0.072 | 0.107 | 0.090 | 0.119 |
| Chr 17 | 0.158 | 0.182 | 0.105 | 0.112 | 0.168 | 0.144 | 0.183 |
| Chr 18 | 0.071 | 0.081 | 0.049 | 0.058 | 0.063 | 0.059 | 0.079 |
| Chr 19 | 0.152 | 0.175 | 0.100 | 0.110 | 0.189 | 0.138 | 0.200 |
| Chr 20 | 0.117 | 0.136 | 0.074 | 0.084 | 0.120 | 0.101 | 0.131 |
| Chr 21 | 0.061 | 0.083 | 0.047 | 0.045 | 0.074 | 0.062 | 0.077 |
| Chr 22 | 0.118 | 0.131 | 0.073 | 0.090 | 0.122 | 0.104 | 0.128 |
| Chr X | 0.029 | 0.039 | 0.019 | 0.023 | 0.041 | 0.030 | 0.034 |
| Chr Y | 0.001 | 0.005 | 0.001 | 0.003 | 0.006 | 0.001 | 0.004 |
| **Overall** | **0.082** | **0.097** | **0.054** | **0.063** | **0.084** | **0.073** | **0.094** |

**Supplementary Table 3.** Mean GS scores for functional elements in the *HBB* gene complex.

|  | Name | Start<sup>a</sup> | Stop<sup>a</sup> | Brain | GI | Lung | Heart | Blood | Muscle | Epithelium |
| --- | --- | --- | --- | --- | --- | --- | --- | --- | --- | --- |
| **Genes in *HBB* gene complex** | *HBB* | 5246696 | 5248301 | 0.0232 | 0.0100 | 0.0017 | 0.0013 | 0.7336 | 0.0008 | 0.0029 |
|  | *HBD* | 5254059 | 5255858 | 0.0013 | 0.0138 | 0.3161 | 0.0014 | 0.8895 | 0.0010 | 0.0340 |
|  | *HBBP1* | 5263184 | 5264822 | 0.0012 | 0.0097 | 0.0527 | 0.0020 | 0.4709 | 0.0008 | 0.0258 |
|  | *HBG1* | 5269502 | 5271087 | 0.0010 | 0.0016 | 0.0017 | 0.0014 | 0.0668 | 0.0008 | 0.0030 |
|  | *HBG2* | 5274421 | 5276011 | 0.0010 | 0.0015 | 0.0018 | 0.0013 | 0.1430 | 0.0008 | 0.0033 |
|  | *HBE1* | 5289580 | 5291373 | 0.0135 | 0.0007 | 0.0017 | 0.0015 | 0.0003 | 0.0008 | 0.0034 |
| **CRMs in *HBB* gene complex** | 3'HS1 | 5226013 | 5226493 | 0.0063 | 0.0089 | 0.0346 | 0.0186 | 0.0090 | 0.0130 | 0.3177 |
|  | HBB_3'enh | 5245876 | 5246140 | 0.0014 | 0.0005 | 0.0022 | 0.0013 | 0.9062 | 0.0008 | 0.0030 |
|  | HBB_prom | 5248301 | 5248556 | 0.0018 | 0.0150 | 0.0016 | 0.0013 | 0.0003 | 0.0008 | 0.0023 |
|  | HBD_prom | 5255713 | 5256160 | 0.0014 | 0.0005 | 0.3528 | 0.0013 | 0.9999 | 0.0008 | 0.0595 |
|  | HBG1_3'enh | 5268365 | 5269114 | 0.0010 | 0.0005 | 0.0193 | 0.0013 | 0.6284 | 0.0008 | 0.0216 |
|  | HBG1_prom | 5271086 | 5271290 | 0.0010 | 0.0005 | 0.0017 | 0.0013 | 0.0003 | 0.0008 | 0.0030 |
|  | HBG1_up | 5271291 | 5271813 | 0.0010 | 0.0005 | 0.0201 | 0.0013 | 0.0494 | 0.0008 | 0.0029 |
|  | HBG2_prom | 5276011 | 5276214 | 0.0010 | 0.0005 | 0.0017 | 0.0013 | 0.0040 | 0.0008 | 0.0030 |
|  | HBG2_up | 5276215 | 5276745 | 0.0010 | 0.0005 | 0.0270 | 0.0013 | 0.0535 | 0.0043 | 0.0204 |
|  | HBE1_prom | 5291175 | 5291343 | 0.0010 | 0.0011 | 0.0017 | 0.0016 | 0.0003 | 0.0008 | 0.0047 |
|  | HBE1_up | 5291344 | 5291610 | 0.0041 | 0.0006 | 0.0017 | 0.0120 | 0.0008 | 0.0008 | 0.0031 |
|  | HBE1_PRB | 5292690 | 5292886 | 0.0010 | 0.0005 | 0.0016 | 0.0021 | 0.0003 | 0.0008 | 0.0015 |
|  | HBE1_NRB | 5292886 | 5292928 | 0.0010 | 0.0005 | 0.0017 | 0.0013 | 0.0003 | 0.0008 | 0.0030 |
|  | HBE1_PRA | 5293982 | 5294081 | 0.0010 | 0.0005 | 0.0017 | 0.0013 | 0.3566 | 0.0008 | 0.0030 |
|  | HBE1_NRA | 5294082 | 5294308 | 0.0010 | 0.0005 | 0.0017 | 0.0013 | 0.0535 | 0.0008 | 0.0030 |
|  | HS1 | 5296894 | 5297517 | 0.0010 | 0.0007 | 0.0403 | 0.0013 | 0.9625 | 0.0040 | 0.1062 |
|  | HS2_pos | 5301795 | 5302089 | 0.2231 | 0.4767 | 0.0604 | 0.4046 | 0.6358 | 0.0336 | 0.7855 |
|  | HS2_neg | 5302090 | 5302174 | 0.0833 | 0.4471 | 0.0635 | 0.0094 | 1.0000 | 0.0051 | 0.9195 |
|  | HS3 | 5305882 | 5306169 | 0.0010 | 0.0005 | 0.0017 | 0.0013 | 0.5671 | 0.0008 | 0.0044 |
|  | HS3.1 | 5306356 | 5306418 | 0.0010 | 0.0005 | 0.0022 | 0.0013 | 0.0233 | 0.0008 | 0.0030 |
|  | HS3.2 | 5306814 | 5307392 | 0.0010 | 0.0005 | 0.0071 | 0.0013 | 0.4680 | 0.0008 | 0.9783 |
|  | HS4 | 5309419 | 5309707 | 0.0010 | 0.2965 | 0.5436 | 0.0013 | 0.4414 | 0.0008 | 0.9378 |
|  | HS5 | 5312534 | 5312694 | 0.0010 | 0.0163 | 0.0543 | 0.0013 | 0.0890 | 0.0008 | 0.8171 |

<sup>a</sup> hg19 coordinates

**Supplementary Table 4.** Mean GS scores for functional elements near *MYH6* and *MYH7.*

| **Name** | **Location<sup>a</sup>** | **Brain** | **GI** | **Lung** | **Heart** | **Blood** | **Muscle** | **Epithelium** |
| --- | --- | --- | --- | --- | --- | --- | --- | --- |
| *MYH6* | chr14: 23851199-23877486 | 0.0562 | 0.0055 | 0.0052 | 0.6050 | 0.0356 | 0.1749 | 0.0833 |
| *MYH7* | chr14: 23881947-23904870 | 0.2315 | 0.0199 | 0.0051 | 0.9788 | 0.0028 | 0.7498 | 0.0739 |
| mir208a | chr14: 23857805-23857875 | 0.0010 | 0.0011 | 0.0026 | 0.2331 | 0.0004 | 0.0040 | 0.2066 |
| mir208b | chr14: 23887196-23887272 | 0.0010 | 0.0011 | 0.0028 | 1.0000 | 0.0004 | 0.4221 | 0.0047 |
| hs1670 | chr14: 23906587-23908214 | 0.9203 | 0.9660 | 0.0306 | 1.0000 | 0.1639 | 1.0000 | 0.2983 |
| hs2330 | chr14: 23911613-23912923 | 0.2340 | 0.2818 | 0.0024 | 0.9999 | 0.0025 | 0.9999 | 0.1892 |

<sup>a</sup> Coordinates are based on hg19.

**Supplementary Table 5.** Cell types used for developing GenoSkyline annotations of ESC and fetal cells.

| Group | EID<sup>a</sup> | Anatomy | Standardized Epigenome Name | Type |
| --- | --- | --- | --- | --- |
| ESC | E002 | ESC | ES-WA7 Cells | PrimaryCulture |
|  | E008 | ESC | H9 Cells | PrimaryCulture |
|  | E001 | ESC | ES-I3 Cells | PrimaryCulture |
|  | E015 | ESC | HUES6 Cells | PrimaryCulture |
|  | E014 | ESC | HUES48 Cells | PrimaryCulture |
|  | E016 | ESC | HUES64 Cells | PrimaryCulture |
|  | E003 | ESC | H1 Cells | PrimaryCulture |
|  | E024 | ESC | ES-UCSF4 Cells | PrimaryCulture |
| Fetal cells | E017 | LUNG | IMR90 fetal lung fibroblasts Cell Line | CellLine |
|  | E093 | THYMUS | Fetal Thymus | PrimaryTissue |
|  | E082 | BRAIN | Fetal Brain Female | PrimaryTissue |
|  | E081 | BRAIN | Fetal Brain Male | PrimaryTissue |
|  | E089 | MUSCLE | Fetal Muscle Trunk | PrimaryTissue |
|  | E090 | MUSCLE_LEG | Fetal Muscle Leg | PrimaryTissue |
|  | E083 | HEART | Fetal Heart | PrimaryTissue |
|  | E092 | GI_STOMACH | Fetal Stomach | PrimaryTissue |
|  | E085 | GI_INTESTINE | Fetal Intestine Small | PrimaryTissue |
|  | E084 | GI_INTESTINE | Fetal Intestine Large | PrimaryTissue |
|  | E086 | KIDNEY | Fetal Kidney | PrimaryTissue |
|  | E088 | LUNG | Fetal Lung | PrimaryTissue |
|  | E080 | ADRENAL | Fetal Adrenal Gland | PrimaryTissue |
|  | E099 | PLACENTA | Placenta Amnion | PrimaryTissue |
|  | E091 | PLACENTA | Placenta | PrimaryTissue |

<sup>a</sup> Epigenome ID in Roadmap project

**Supplementary Table 6.** List of 15 complex diseases and traits.

| Trait/Disease | Category | Data source | Data Link | Ref. |
| --- | --- | --- | --- | --- |
| Schizophrenia | Psychiatric disease | PGC | http://www.med.unc.edu/pgc/downloads | 31 |
| Anorexia nervosa | Psychiatric disease | GCAN | http://www.med.unc.edu/pgc/files/resultfiles/gcan_meta-out.gz | 44 |
| Coronary artery disease | Cardiovascular disease | CARDIoGRAM | http://www.cardiogramplusc4d.org/downloads/ | 33 |
| Crohn's disease | IBD | IIBDGC | http://www.ibdgenetics.org/downloads.html | 45 |
| Ulcerative colitis | IBD | IIBDGC | http://www.ibdgenetics.org/downloads.html | 46 |
| Rheumatoid arthritis | Inflammatory disorder | Stahl et al | http://www.broadinstitute.org/ftp/pub/rheumatoid_arthritis/Stahl_etal_2010NG/ | 47 |
| LDL cholesterol | Lipids | Global lipids consortium | http://csg.sph.umich.edu/abecasis/public/lipids2013/ | 48 |
| HDL cholesterol | Lipids | Global lipids consortium | http://csg.sph.umich.edu/abecasis/public/lipids2013/ | 48 |
| Triglycerides | Lipids | Global lipids consortium | http://csg.sph.umich.edu/abecasis/public/lipids2013/ | 48 |
| Total cholesterol | Lipids | Global lipids consortium | http://csg.sph.umich.edu/abecasis/public/lipids2013/ | 48 |
| Type 2 diabetes | Metabolic disorder | DIAGRAM | http://diagram-consortium.org/downloads.html | 49 |
| Bone mineral density | Osteoporosis | GEFOS | http://www.gefos.org/?q=content/data-release | 50 |
| BMI | Anthropometric traits | GIANT | http://www.broadinstitute.org/collaboration/giant/index.php | 51 |
| WHR adjusted for BMI | Anthropometric traits | GIANT | http://www.broadinstitute.org/collaboration/giant/index.php | 29 |
| Height | Anthropometric traits | GIANT | http://www.broadinstitute.org/collaboration/giant/index.php | 52 |

**Supplementary Table 7.**
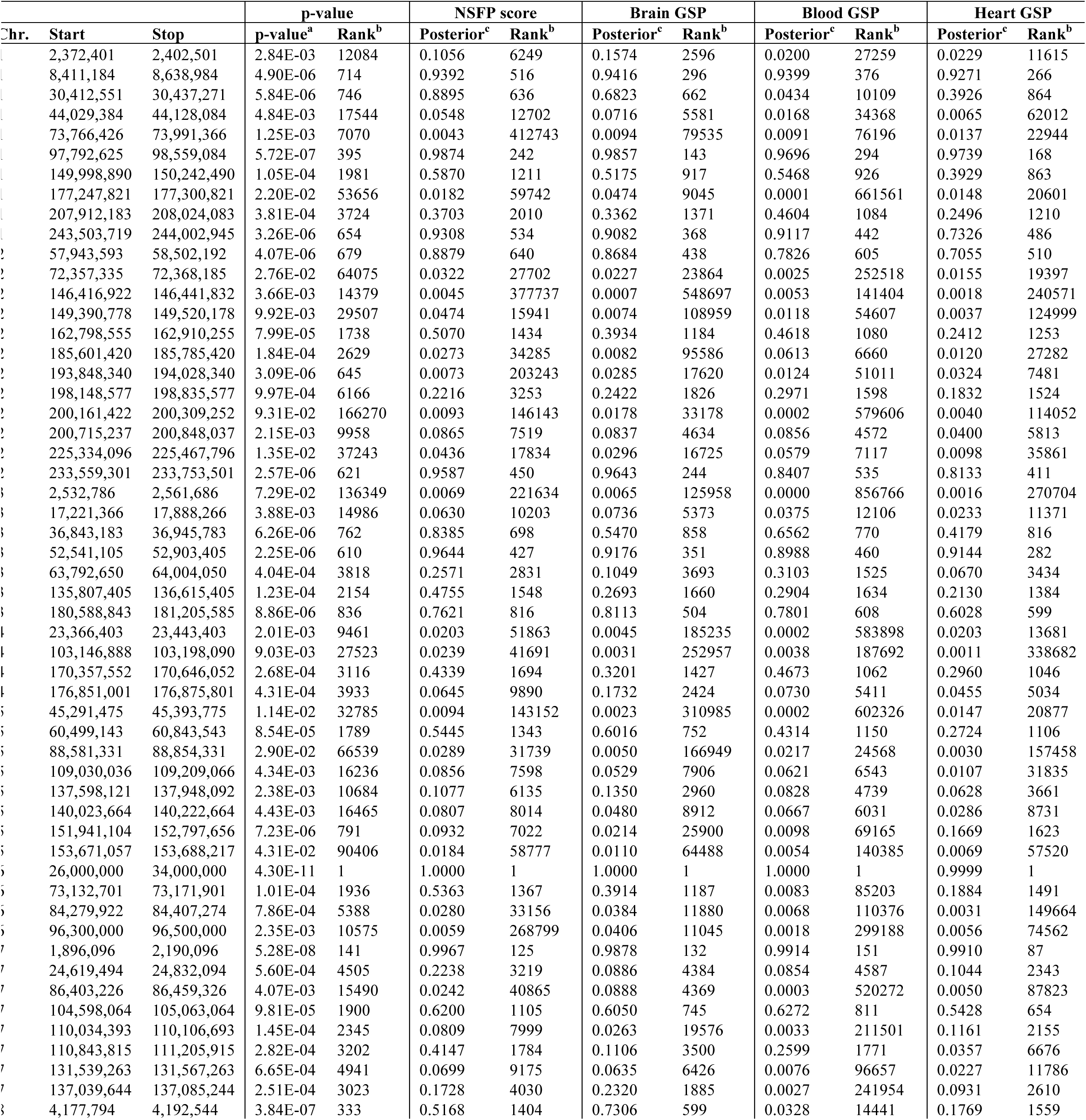
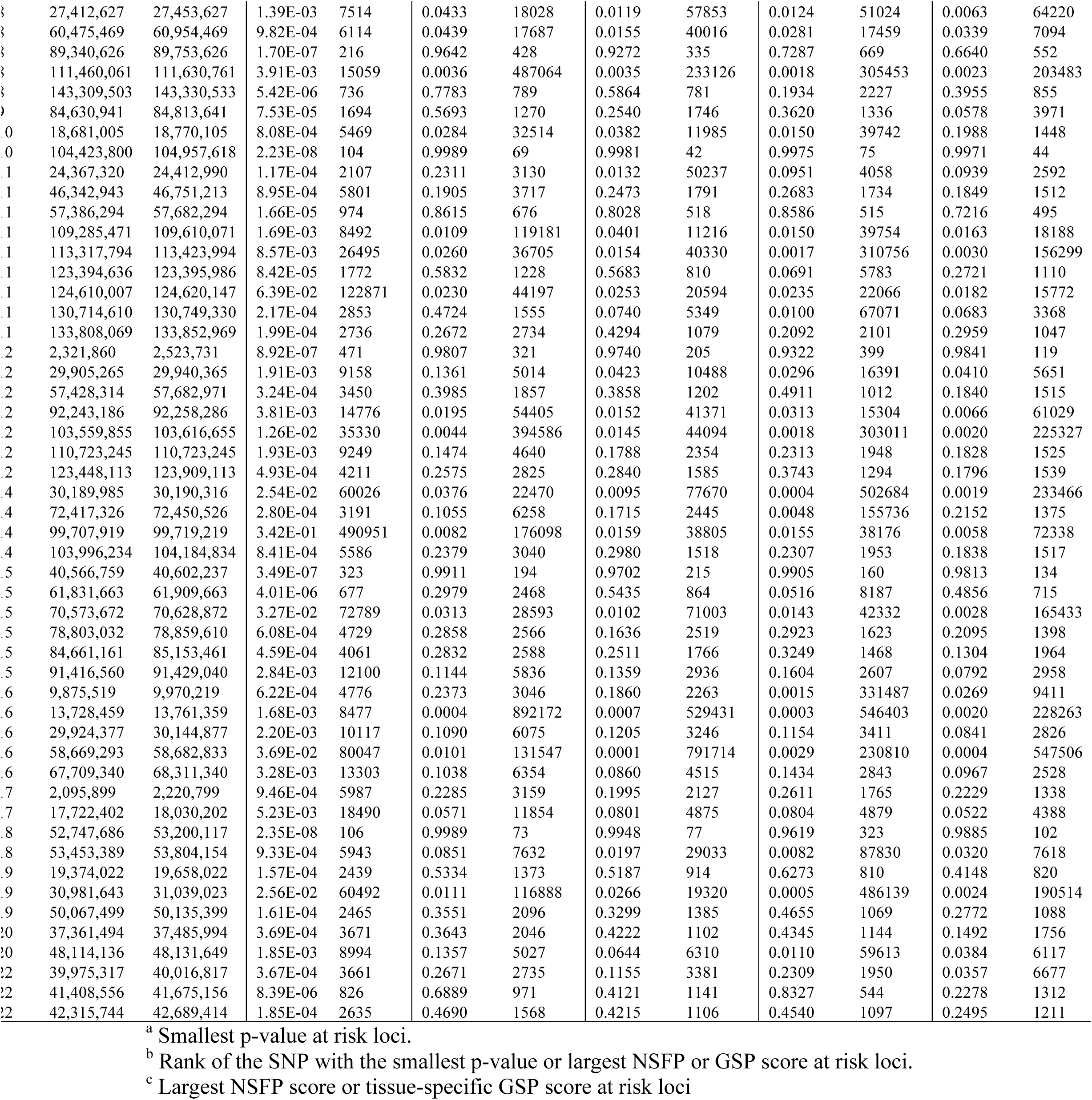
Ranks of top signals under different criteria at 105 schizophrenia-associated loci.

**Supplementary Table 8.**
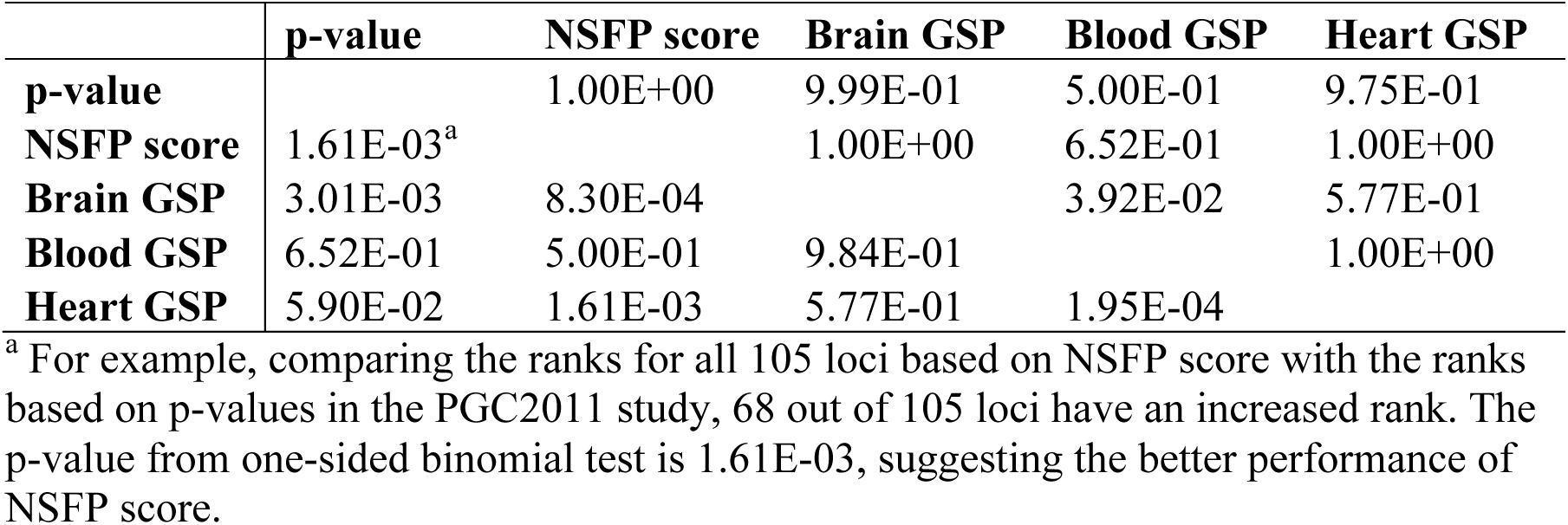
Ranking performance comparison. The value in each cell is the p-value acquired from one-sided binomial test.

|  | **p-value** | **NSFP score** | **Brain GSP** | **Blood GSP** | **Heart GSP** |
| --- | --- | --- | --- | --- | --- |
| **p-value** |  | 1.00E+00 | 9.99E-01 | 5.00E-01 | 9.75E-01 |
| **NSFP score** | 1.61E-03<sup>a</sup> |  | 1.00E+00 | 6.52E-01 | 1.00E+00 |
| **Brain GSP** | 3.01E-03 | 8.30E-04 |  | 3.92E-02 | 5.77E-01 |
| **Blood GSP** | 6.52E-01 | 5.00E-01 | 9.84E-01 |  | 1.00E+00 |
| **Heart GSP** | 5.90E-02 | 1.61E-03 | 5.77E-01 | 1.95E-04 |  |

<sup>a</sup> For example, comparing the ranks for all 105 loci based on NSFP score with the ranks based on p-values in the PGC2011 study, 68 out of 105 loci have an increased rank. The p-value from one-sided binomial test is 1.61E-03, suggesting the better performance of NSFP score.

**Supplementary Table 9.**
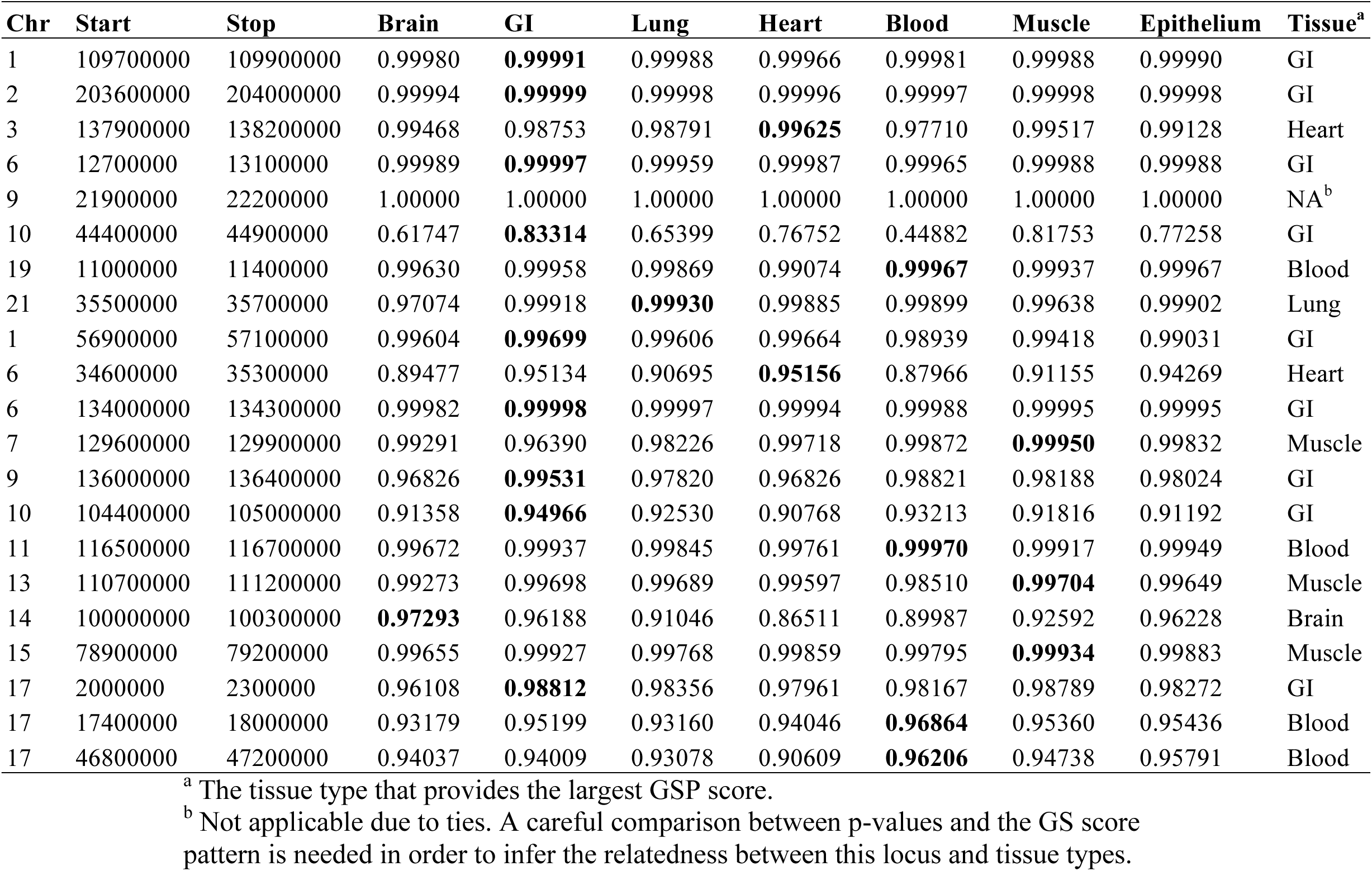
Largest GSP scores for each tissue type at CAD-associated risk loci.

**Supplementary Table 10.**
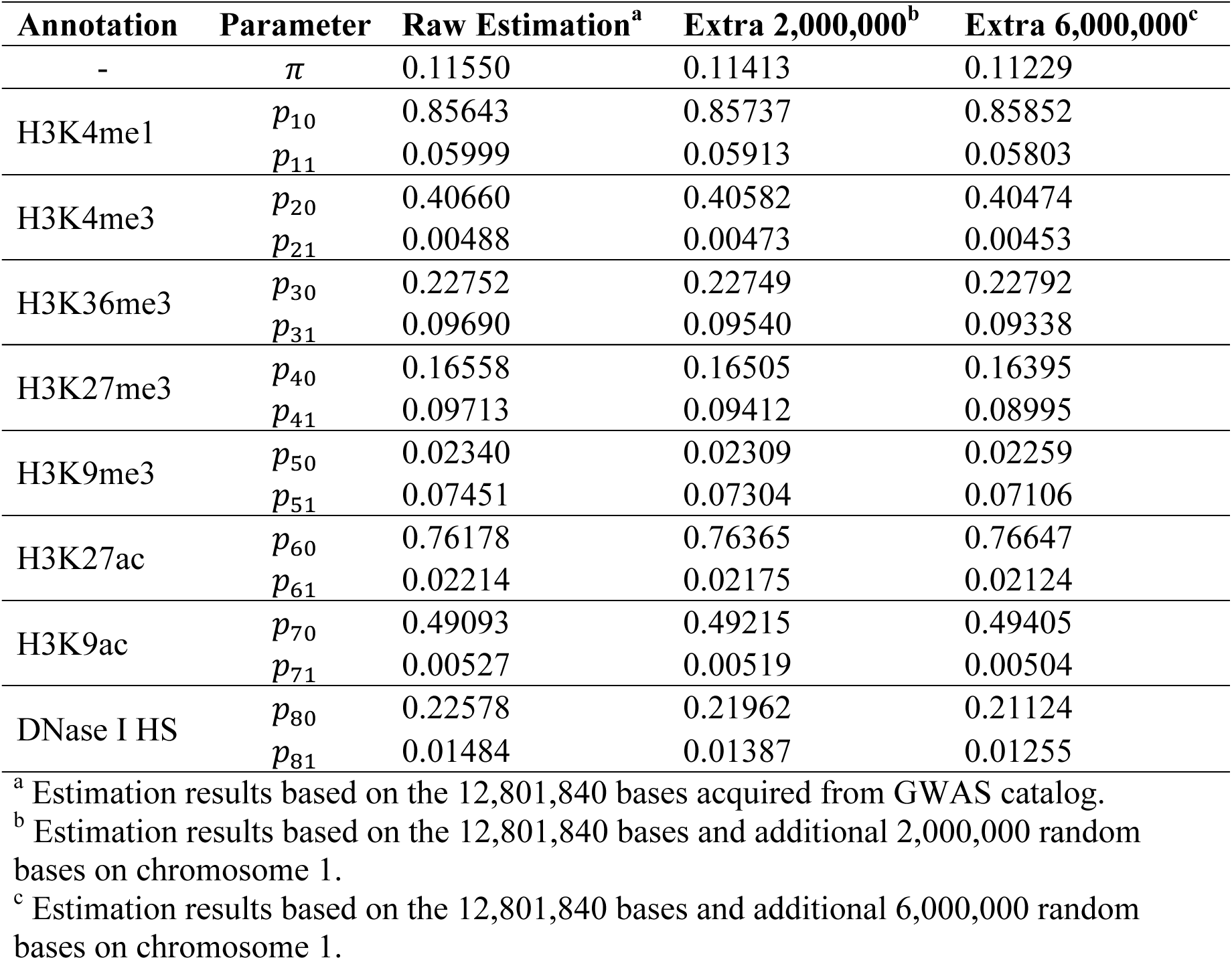
GenoSkyline parameter estimates for brain tissue.

